# SpliceDecoder: A High-Throughput Tool for Guiding the Functional Interpretation of Differential Splicing Events

**DOI:** 10.1101/2025.10.01.679902

**Authors:** Hyeon Gu Kang, Marina Yurieva, Mattia Brugiolo, Jeffrey Chuang, Olga Anczukow

**Affiliations:** The Jackson Laboratory for Genomic Medicine, Farmington, CT, USA; Department of Genetics and Genome Sciences, UConn Health, Farmington, CT, USA; Institute for Systems Genomics, UConn Health, Farmington, CT, USA

## Abstract

Alternative splicing generates different mRNA isoforms from single genes, generating protein diversity essential for normal development and tissue function. Dysregulation of splicing is implicated in numerous diseases, including cancer, neurodegeneration, diabetes, and rare genetic disorders. Although thousands of spliced isoforms have been identified in disease-relevant contexts, the functional significance of most remains unknown, hindering our understanding of splicing-driven disease mechanisms and limiting therapeutic discovery. A major challenge lies in prioritizing biologically meaningful splicing events identified through short-read RNA sequencing. Existing splicing analysis tools typically rank target events based on splicing change magnitude or gene-level annotation, often without evaluating how resulting isoforms impact protein structure or function. This creates a critical bottleneck in translating splicing data into biological and therapeutic insights. To address this, we developed SpliceDecoder, a computational workflow that predicts how each splicing event or isoform impacts transcript productivity, protein sequence, and functional domains. Each event is assigned a functional effect score to guide evidence-based prioritization. SpliceDecoder facilitates a more informed interpretation of splicing data, reduces reliance on prior knowledge, and enables identification of events with potential biological and clinical relevance. We demonstrate its utility by validating known splicing alterations and identifying novel disease-associated isoform switches across public datasets.

## Introduction

Alternative splicing is a complex post-transcriptional regulatory process that allows a single gene to produce multiple mRNA isoforms, thereby expanding the coding potential of the genome and contributing to the functional complexity of higher organisms (1). By selectively including or excluding exons or introns, alternative splicing can influence transcript stability, subcellular localization, and the structure and function of encoded proteins. This process is tightly regulated in a cell-type, tissue-specific, and developmental context, and is essential for normal physiology, including processes such as cell differentiation, signaling, and immune response (2,3). Disruptions in splicing regulation have been increasingly recognized as a hallmark of human disease (4–6). Advances in high-throughput RNA-sequencing have revealed widespread changes in splicing patterns in disease-relevant tissues and conditions, including in cancer, neurodegenerative disorders, metabolic syndromes, and rare genetic diseases. For example, in cancer, a splicing switch from the pyruvate kinase *PKM1* to the *PKM2* isoform promotes aerobic glycolysis and tumor growth (7,8), whereas splicing of the cell death regulator *BCL2* generates isoforms with opposing roles in apoptosis, influencing cancer cell survival and chemoresistance (9–11). Similarly, a splicing switch in the *HRAS* oncogene creates either a full-length p21 protein or a truncated p19 protein, which lacks 16 amino acids in the C-terminal domain and differs in its ability to bind GTP (12–14). Finally, splicing of myosin phosphatase RHO-interacting protein *MPRIP* is associated with metastasis, altered cytoskeletal organization and formation of focal adhesions in pancreatic cancer (15). Importantly, therapies that modulate splicing of *PKM*, *BCL2*, *HRAS,* or *MPRIP* isoforms have shown efficacy in pre-clinical models (11,12,15–19) demonstrating that spliced isoforms represent clinically actionable and successful targets for human disease applications. These examples illustrate how changes in splicing patterns can drive disease by affecting the expression, structure, and function of key proteins, highlighting the need to better understand the functional consequences of spliced isoforms in a more systematic manner.

However, despite significant advances in detecting differential splicing (DS) using short-read RNA sequencing (RNA-seq), distinguishing functionally relevant splicing events from background noise remains a major challenge. This in turn impedes the discovery of clinically relevant and therapeutically actionable splicing alterations. A variety of computational tools have been developed to identify DS event in pre-clinical and clinical RNA-seq datasets (20), which generally fall into two broad categories. Event-centric approaches – such as rMATS (21) – quantify changes in exon or splice junction usage between conditions, and, transcript-centric approaches – such as Stringtie (22) – aim to reconstruct and quantify full-length transcripts from short reads. Both strategies offer complementary insights but also face limitations in accurately linking splicing changes to functional outcomes. Event-centric tools are particularly popular due to their robustness and statistical power, often reporting percent spliced-in (PSI) values and false discovery rates (FDR) for thousands of splicing events (20). However, these methods typically lack the capacity to assess how each splicing event may affect the structure or function of the encoded protein. Functional interpretation requires knowledge of the impacted transcript isoforms and the functional domains they encode—information that is not readily accessible from standard event-centric outputs. Thus, it remains challenging to sort through and curate a list of the most likely biologically meaningful differentially spliced events (DSEs) from the thousands of statistically significant events identified by these tools. As a result, researchers frequently prioritize DSEs based on PSI or FDR ranking or prior gene-level knowledge, potentially overlooking biologically meaningful but less obvious isoforms. This reliance on *a priori* knowledge and statistical thresholds introduces bias and limits the translation of splicing data into mechanistic insight and therapeutic opportunities.

Here, we develop SpliceDecoder, a high-throughput, Python-based tool designed to facilitate the functional interpretation of DSEs or isoforms identified using event- or transcript-centric RNA-seq analyses. SpliceDecoder seamlessly integrates with the output of widely used DS analysis pipelines (23), and enables researchers to rank DSEs based on their predicted functional consequences rather than solely on statistical significance. The workflow comprises three main steps: 1) a preprocessing step that reformats event-based splicing data, 2) a mapping step that links each DSE to transcript models sharing the exact exon structure, and 3) a simulation step that compares simulated and reference transcripts to infer predicted functional effects. SpliceDecoder evaluates and scores predicted outcomes such as the introduction of premature stop codons and likelihood of nonsense mediated decay (NMD), disruption a protein domain, or alterations in untranslated regions (UTRs). Additionally, it outputs FASTA-formatted amino acid sequences for both the reference and simulated transcripts, enabling downstream structural modelling with AI-based tools such as AlphaFold2 (24). Notably, we find that SpliceDecoder functional impact scores are often not correlated with traditional metrics such as PSI or FDR, indicating that statistical significance alone is not a reliable predictor of biological relevance. To address this, we propose a combinatorial ranking strategy that integrates SpliceDecoder’s functional predictions with splicing magnitude and statistical significance to more effectively prioritize biological meaningful DSEs. We demonstrate the utility of this approach by validating known functional events and identifying novel isoform switches across publicly available datasets.

## Materials and Methods

### Event-based splicing analysis using RNA-seq data

We used previously described RNA-seq from human mammary epithelial MCF-10A cells at 0, 8, and 24 hours after MYC activation (n=3; GSE181968) (14). RNA-seq reads were aligned to human reference genome using STAR (v.2.7.3a) (25) in 2-pass mode and differential splicing (DS) analysis was performed using rMATS (v.4.0.2) (21) comparing MYC activation at 0h *vs.* 8h and 0h *vs.* 24h using our existing pipeline https://github.com/TheJacksonLaboratory/splicing-pipelines-nf (**Supplementary Table 1** and **2**). From the ‘JCEC.txt’ rMATS output files, we retained splicing events with ≥ 5 junction reads and FDR ≤ 0.05 and |IncLevelDifference| ≤ 0.1. Genomic coordinates of each exon were defined as the positions of the ‘upstream ES/EE’, ‘downstream ES/EE’, and ‘exonStart_0base’ and ‘exonEnd’ or ‘riExonStart_0base’ and ‘riExonEnd’, ‘longExonStart_0base’ and ‘longExonEnd’. Finally, we defined *ΔPSI* as ‘group2 (MYC-active) PSI – group1(MYC-inactive) PSI’. This pre-processing step for downstream analysis was performed by applying a custom python script ‘make_input_from_rmats.py’ to the JCEC files from the rMATS output and the processed data was used as the test dataset.

### Mapping splicing events to the whole transcriptome

For each splicing event, SpliceDecoder generates all possible splicing cases for each event type (CA-Cassette Alternative exon, RI-Retained Intron, A3SS-Alternative 3’ Splice Site, A5SS-Alternative 5’ Splice Site, and MXE-Mutually Exclusive Exons) (**Supplementary Fig. 1A**). For example, exon skipping (ES) and inclusion (EI) cases are generated from each CA event. SpliceDecoder uses genomic coordinates of each DSE as specific target positions for each event type: (1) E1 end/E2 start/E2 end/E3 start positions are considered for exon inclusion (EI), alternative 5’/3’ splice site (Alt_A5/3SS), and mutually exclusive exon 1 and 2 (MXE1/2); (2) E1 end/E3 start positions are considered for exon skipping (ES), skipped intron (SI), canonical 5’/3’ splice site (Can_A5/3SS); (3) E1 start/E3 end positions for retained intron (RI).

Then, SpliceDecoder uses the specified target positions to extract isoforms, from the GTF file provided to rMATS. To identify isoforms with exon structures that perfectly match the target positions of each DSE, SpliceDecoder performs a tiling-based comparison. This involves sequentially aligning the target positions of each DSE with the exon tiles of transcripts, starting from the first exon of all isoforms within the same gene (**Supplementary Fig. 1B**). SpliceDecoder compares the start and end genomic coordinates of each exon tiles and excluded transcripts with even a single base pair mismatch. To avoid false matches in Alt_A5SS and A3SS, caused by intervening exons between E1 and E3, we apply additional filtering criteria. For Alt_A3SS, the exon aligned with E2 start must be after E2 end. For Alt_A5SS, the exon aligned with E2 end must be before E2 start (**Supplementary Fig. 1B**). Isoforms that match all target positions without mismatches are defined as Ref-TX for each splicing case. Although relaxing a mapping threshold (*e.g.*, allowing >1 base difference) could increase the mapping rate, even one base difference may alter the reading frame. Therefore, only perfectly matched transcripts are used in downstream analyses.

To evaluate the accuracy of the mapping process, we examined DSEs that failed to map. Specifically, we examined unmapped DSEs that resulted from mapping error or represent novel events predicted by rMATS. We utilized the *novelJunction.txt* file generated by rMTAS, which lists junctions not present in the given GTF file. All unmapped DSEs were confirmed to correspond to novel events identified by rMATS.

### Simulating alternatively spliced transcripts

To estimate the functional impact of DSEs, SpliceDecoder generates simulated transcripts (Sim-TXs) by modifying the exon structure of the matched Ref-TX to reflect the corresponding alternative splicing pattern. Each Sim-TX retains the exon structure of Ref-TX, except in the region affected by alternative splicing. For example, to simulate exon inclusion (*simEI*), the Sim-TX is created by inserting the skipped exon into the Ref-TX that corresponds to the skipped exon structure (**Fig. 1A**). Conversely, to simulate exon skipping (*simES*), the Sim-TX is generated by removing the included exon from the Ref-TX that corresponds to the included exon structure. The same strategy is applied to simulate retained/skipped intron (*simRI/SI*), alternative/canonical 3’ and 5’ splice site Sim-TX (*simAlt/Can_A3/5SS*), and mutually exclusive exon (*simMXE1/2*). All simulation steps generate Ref- and Sim-TX BED and FASTA files for further processing steps.

**Figure 1.**
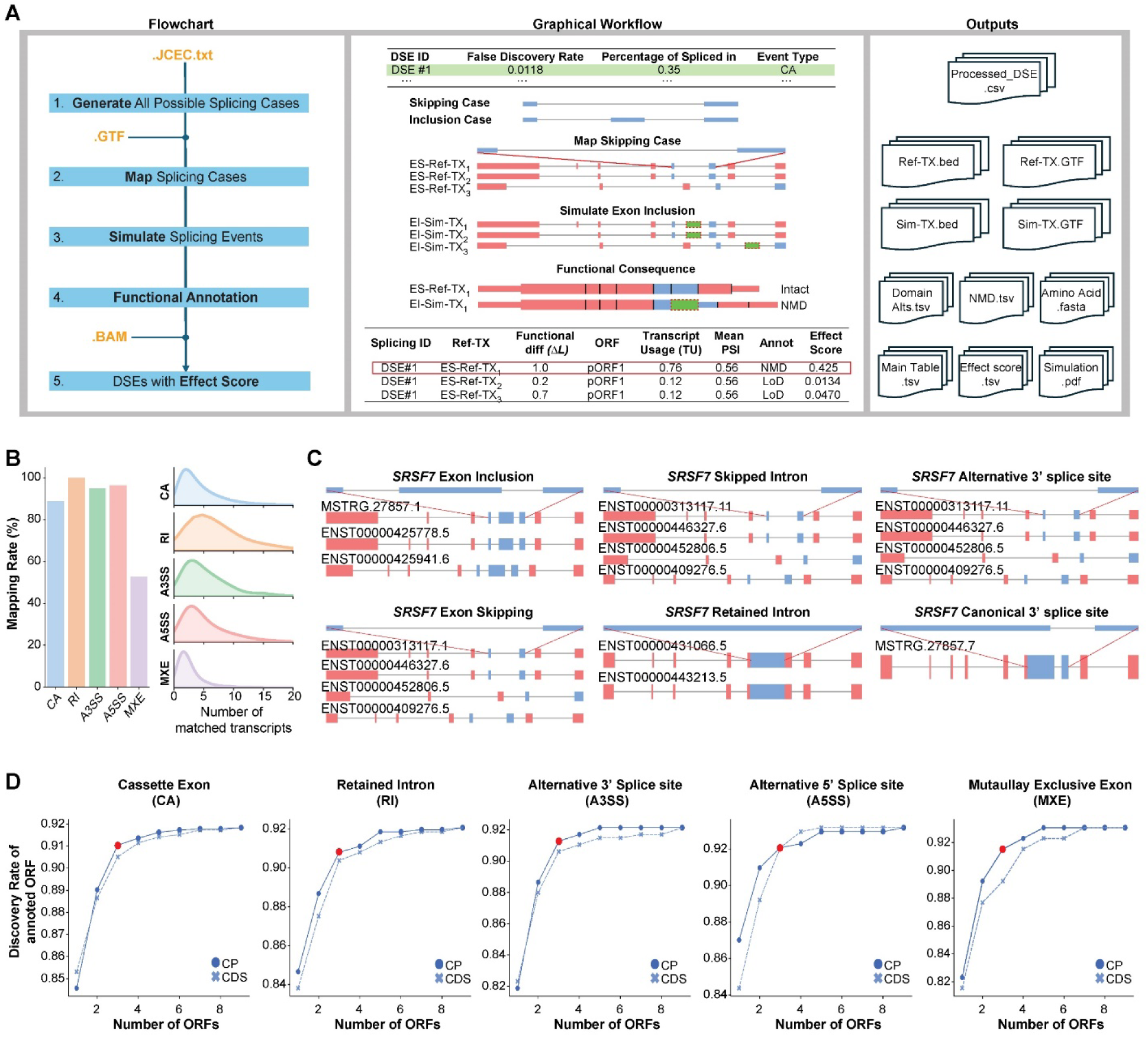
Overview of the SpliceDecoder workflow. (**A**) Schematic of SpliceDecoder workflow displaying the core steps of SpliceDecoder and their input file formats in orange (left). A Graphical workflow box provides an example using a cassette exon (CA) event (middle). An output box introduces the SpliceDecoder output files generated at each step of the Flowchart (right). (**B**) Mapping rates of DSEs (left) and number of matched transcripts for query cases (right) shown per splicing event type; cassette exon (CA), retained intron (RI), alternative 3’/5’ splice site (A3/5SS), and mutually exclusive exon (MXE), for our test dataset comparing human mammary epithelial MCF-10A cells at 8h after activation of the MYC oncogene vs. control cells. (**C**) Examples of mapped query cases with their matched transcripts, with the gene name and type of query case indicated above and the transcript structures depicted below. Matched transcripts are depicted below in red with exons that perfectly match with the query case highlighted with a block line. Each transcript has their own ID that is matched with the given GTF file. (**D**) Discovery rate of annotated ORF (y-axis) according to the number of predicted ORFs (x-axis) based on coding potential (CP) and coding sequence length (CDS). CP and CDS are depicted by circles and ‘x’ marks, with the elbow points indicated by red circles.

### Identifying open reading frames in reference and simulated transcripts

To assign the functional features and identify potential nonsense-mediated decay (NMD) targets, SpliceDecoder predicts all possible open reading frame (ORF) for each Ref-TX and Sim-TX using CPAT2 (26) (https://rna-cpat.sourceforge.net). This analysis uses FASTA files of Ref-TX and Sim-TX genomic sequences as input, along with the Human.v32.logit.RData.

In the absence of reference annotations, we evaluated two criteria for selecting predicted ORFs: (1) coding potential score and (2) coding sequence (CDS) length. For each transcript, we selected the top 9 ORFs based on each criterion and assessed their ability to recover known ORFs annotated in GENCODE. We defined the ORF discovery rate as the proportion of annotated ORFs captured by predicted set, using GENCODE as a reference. By incrementally increasing the number of predicted ORFs from 1 to 9, we compared corresponding discovery rates and identified the elbow point method —beyond which additional ORFs yielded minimal gains. This analysis showed that selecting the top 3 predicted ORF (pORF) was sufficient to recover over 90% of annotated ORFs. Based on this result, SpliceDecoder uses the top 3 pORFs as the default set for functional feature assignment in each DSE (**Fig. 1D**).

### Mapping annotated functional features of reference and simulated transcripts

For each transcript, SpliceDecoder retrieves known protein features using the genomic coordinates of annotated protein domain, protein motif, protein region, DNA binding protein regions, coiled-coil regions, chain, disordered regions, and other structural features from UniProt DB FTP (https://ftp.uniprot.org/pub/databases/uniprot/current_release/knowledgebase/genome_annotation_tracks/) (27). These functional features are assigned to Ref-TX and Sim-TX using the *bedtools intersec*t function (28). To ensure relevance to protein-coding regions, functional features located within untranslated regions (UTRs) are excluded based on the predicted reading frame information.

### Estimating and classifying the functional consequence of alternative splicing

SpliceDecoder uses four sequential steps to compare the functional features between the Ref-TX and Sim-TX within the same reading frame (**Fig. 2A**) and classifies each DSE event into one of the seven functional classes detailed below (LoF, NMD, PTC-removal, LoD, GoD, CDS alt, or UTR alt):

**Figure 2.**
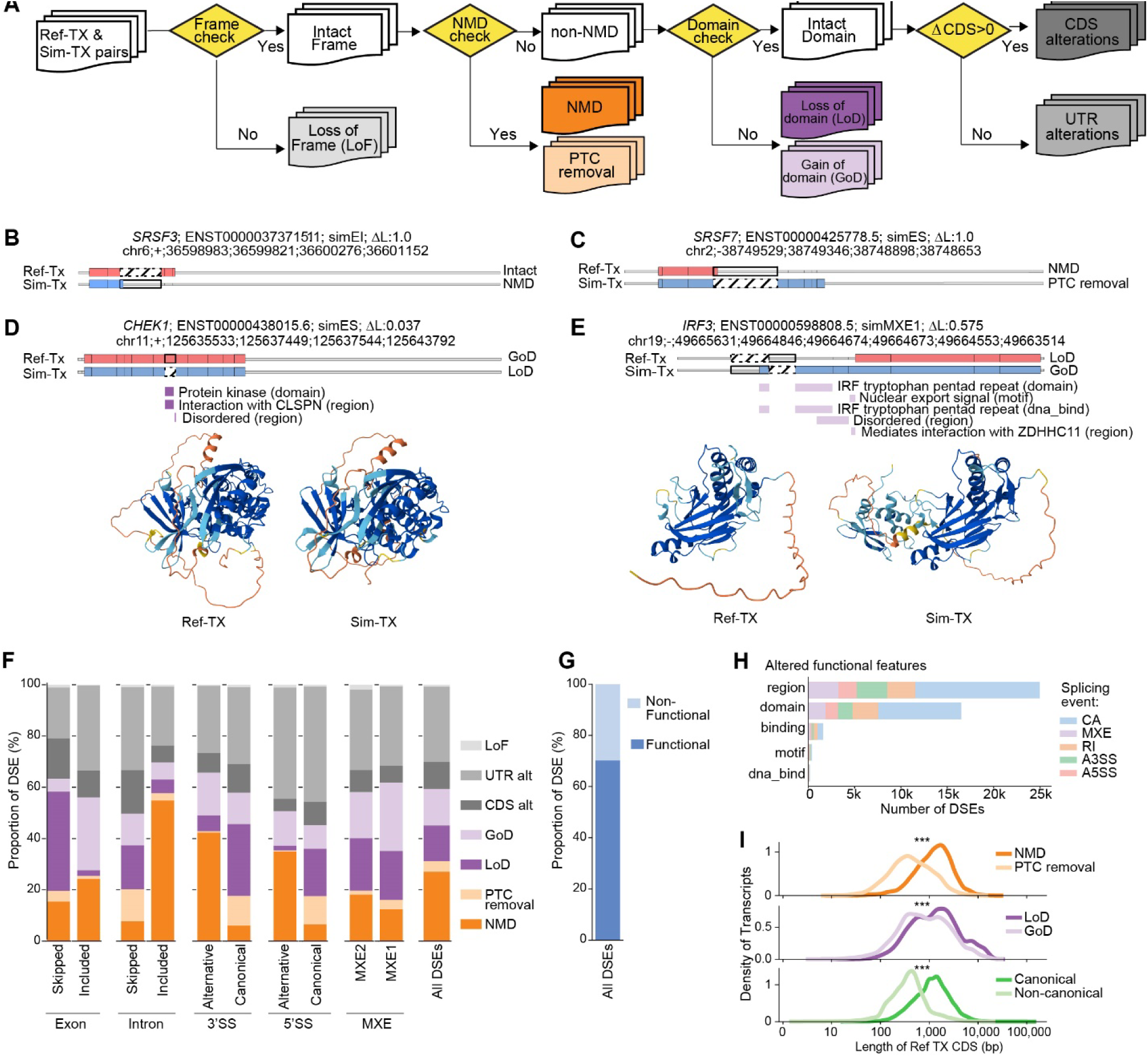
SpliceDecoder enables classifying DSEs by their functional consequence and reveals differences between splicing events. (**A**) Overview of the SpliceDecoder functional class annotation process which predicts functional differences between a simulated transcript (Sim-TX) and its matched transcript (Ref-TX), classifying them into one of seven functional classes: NMD, PTC removal, Gain of Domain (GoD), Loss of Domain (LoD), CDS alterations (CDS alt), UTR alterations (UTR alt), or Loss of Frame (LoF). Cases with domain alterations have a ⊿L value corresponding to the magnitude of their protein domain change, and NMD cases are assigned the maximal value (1). (**B, C**) Examples of NMD (**B**) and PTC removal (**C**) classes. Gene symbol, Ref-TX ID, simulated event, ⊿L, and LongID of rMATS are shown above each simulation. Transcript and exon structures of Ref-TX and Sim-TX are shown below the titles, with UTRs (gray), CDS (pink and blue) regions, and inclusion (bold) or skipping (dotted) of exonic or intronic sequences. Functional classes (NMD, PTC removal, LoD, and GoD) are displayed at the right end of Sim-TX. (**D, E**) Examples of LoD (**D**) and GoD (**E**) classes formatted as in B (**B**). For domain alteration, the location of lost domains are shown below the transcript structure in dark purple (**D**) and gained domains in light purple (**E**), with domain names shown next to each domain. The 3D protein models at the bottom are predicted by AlphaFold2 using the translated amino acid sequences of Ref-TX and Sim-TX provided by SpliceDecoder. (**F**) Distribution of each functional class per splicing event type (left) and for all DSEs (right) detected in human mammary epithelial MCF-10A cells at 8h after MYC activation vs. control. (**G**) Proportion of DSEs with at least one functional consequence (including LoD, GoD, NMD, and PTC removal). (**H**) Distribution of altered functional features across splicing event types. The y-axis represents type of Uniprot functional feature types, and colors indicate different splicing types. Altered domain information was obtained from the Domain_Alts.tsv. (**I**) Length of the matched transcript (Ref-TX) shown by functional consequences, NMD-related, PTC removal, LoD, and GoD, all classes were depicted same color with (**F**), and isoform types (GENCODE canonical and non-canonical). Statistical differences were calculated by 2 sample KS test.

**1)** ‘*Frame Check*’: this step assesses whether the Sim-TX retains the start or stop codon of the Ref-TX. If both codons are preserved, the Sim-TX is labelled as *Intact Frame* and proceeds to the next step. If both codons are absent, the Sim-TXs is labelled as *Loss of Frame* (LoF) and excluded from the downstream analyses.

**2)** ‘*NMD Check*’: this step investigates whether the Sim-TX is likely subject to NMD using the 50-55 nt NMD rule as a basic criterion (29). If a premature stop codon (PTC) is introduced more than 55bp upstream of the last exon-exon junction site, the Sim-TX is classified as *NMD*. Applying the same rule, if a PTC present in the Ref-TX is absent in the Sim-TX, the Sim-TX is classified as *PTC removal*. Sim-TXs with no qualifying PTC changes are classified as *non-NMD* and tested in the next step for possible domain changes.

To complement this basic NMD classification and to identify DSEs that potentially escape NMD, SpliceDecoder provides an advanced criterion where PTC located less than 150 nt downstream of the start codon (Start-proximal) or within an exon longer than 407 nt (Long-exon) are classified as *non-NMD* (30).

**3)** ‘*Domain Check*’: the step investigates whether protein domain and functional features differ between Sim-TX and Ref-TX. SpliceDecoder evaluates individual domain blocks (rather than entire domains) to detect partial gains or losses due to DSEs. For each domain block *i*, a relative functional change ratio (Δ*L_i_*) is calculated as:

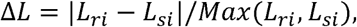

where *L_ri_* and *L_si_* represent the domain length of Ref-TX and Sim-TX respectively. SpliceDecoder calculates the absolute difference |*L_ri_* - *L_si_* | in domain blocks mapped to each exon and normalized it by the maximum domain length Max(*L_ri_, L_si_*). The final domain change ratio, ΔL, is the average of ΔL across all domain blocks. A value of Δ*L=*1 indicates complete domain alteration and is assigned by default to all *NMD* and *PTC removal* events. A value of Δ*L=0* indicates DSE events that do not impact known protein domain and are classified as *Intact Domain*. If Δ*L* ≠ *0*, domain changes are further classified as: *Loss of Domain* (LoD) for Sim-TX with shorter domains, *Gain of Domain* (GoD) for Sim-TX with longer domains compared to Ref-TX. If both GoD and LoD occur, the category with the greater change is assigned. Sim-TX classified as *Intact domain* proceed to the next step.

**4)** ‘*CDS length*’: the step compares the coding sequence (CDS) length of Ref-TX and Sim-TX using the relative CDS change ratio (Δ*CDS*):

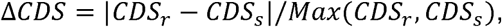

where CDS_r_ and CDS_s_ represent the coding sequence lengths of Ref-TX and Sim-TX respectively. A value of Δ*CDS=*1 indicates complete CDS change and is assigned to all *NMD* and *PTC removal*. For Sim-TX with *Intact Domain*: if Δ*CDS*>0 and Δ*L*=0, the Sim-TX is classified as *CDS alteration* (CDS alt); if Δ*CDS*=0 and Δ*L*=0 the Sim-TX is classified as *UTR alteration* (UTR alt).

### Assessing functional differences between transcripts

To evaluate the functional differences of transcripts identified through a transcript-centric approach, SpliceDecoder categorized transcripts as canonical (Cano-Tx) and non-canonical (Query-Tx) by using the GENCODE GTF. To do this, we used the attribute column of the GENCODE GTF as follows: 1) transcripts tagged as “basic” or “Appris_principal” and lacking the “Could not be confirmed start/end of mRNA or CDS” tag in the attributes, were classified as Cano-Tx; 2) all other transcripts were classified as Query-Tx. Alternatively, users can define Cano-Tx manually by providing a custom list of Cano-Tx and their corresponding gene IDs, allowing flexible classification based on experimental or context-specific conditions.

Then, SpliceDecoder compared each query transcript (Query-Tx) with a corresponding canonical transcript (Cano-Tx) of the same gene. Functional domain mapping was performed based on the ORF with the highest coding potential in each transcript, using the same algorithm that was applied in the event-centric analyses. Functional classification of the Query-Tx was determined by comparing its functional domains and NMD possibility with those of its Cano-Tx, using the same algorithm as in the event-based approach.

### Stratification of canonical and non-canonical isoforms

To define canonical and non-canonical isoforms, we applied the same categorization method to the GENCODE GTF. We considered Cano-Tx as canonical isoforms and Query-Tx as non-canonical isoforms. We compiled these classifications into a transcript dictionary containing transcript IDs and class labels (canonical or non-canonical). CDS lengths were retrieved from Protein-coding transcripts in GENCODE v32 (https://www.gencodegenes.org/human/release_32.html), and integrated into the dictionary file, along with previously assigned functional classes from above.

### Validating NMD predictions from SpliceDecoder

To validate NMD-sensitive Sim-TX predicted by SpliceDecoder, we compared their intron chain structures against GENCODE v32 annotation using GFFcompare. In the resulting ‘refmap’ file, we identified 488 Sim-TXs with a ‘=’ tag, indicating perfect matches to annotated transcripts. Next, we examined the transcript types of these matched Sim-TXs (**Fig. 3A**).

**Figure 3.**
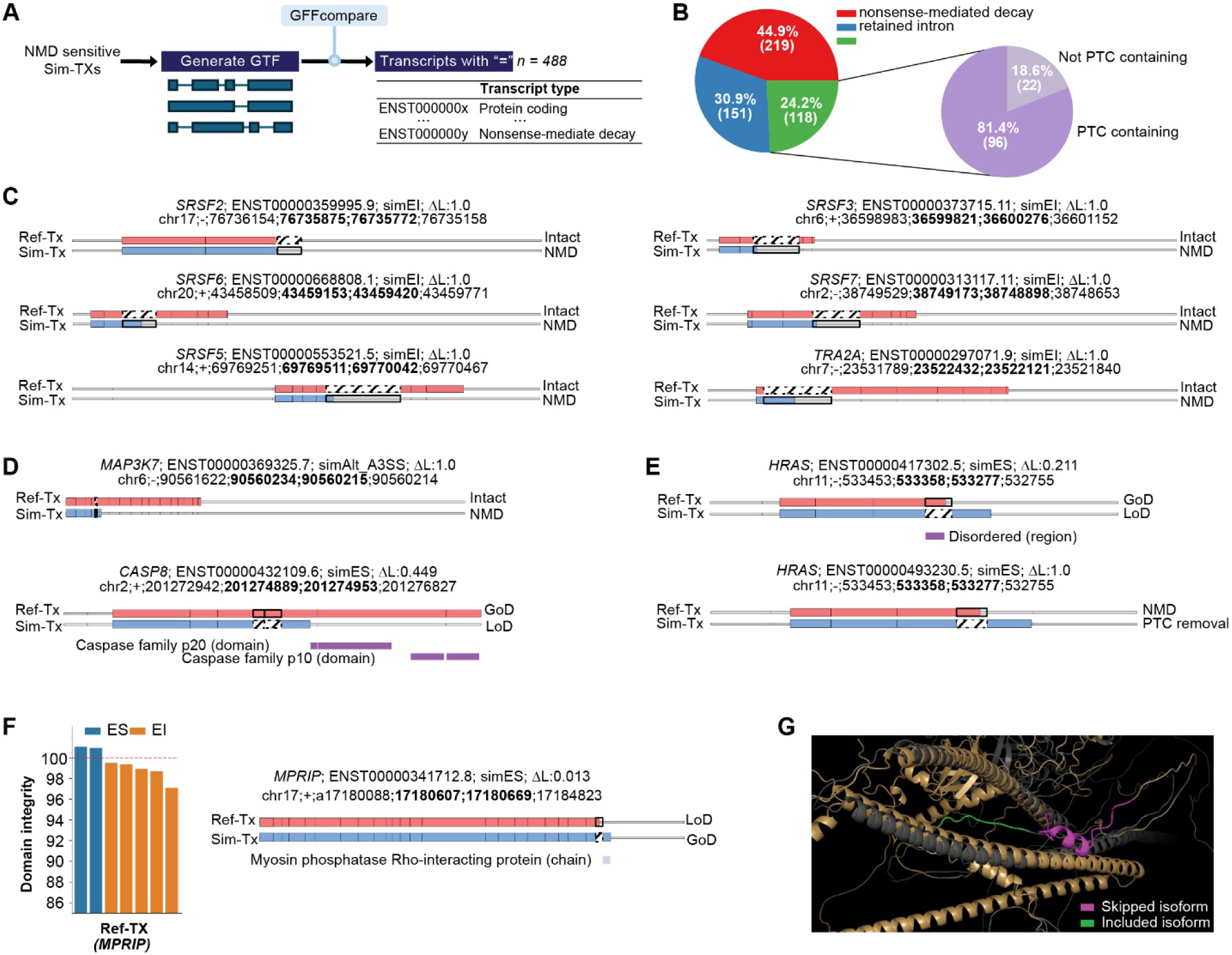
Validation of predicted functional consequences. (**A,B**) Overview of GENCODE based validation for NMD sensitive Sim-TXs (**A**), with the resulting distribution of transcript types for NMD sensitive Sim-TXs shown in (**B**). (**C**) Predicted consequences of experimentally validated poison exons in splicing factors. Each event is annotated with Gene symbol, Ref-TX ID, simulated event and ⊿L is shown for each event along with the transcript and exon structures of Ref-TX and Sim-TX with UTRs (gray), CDS (pink and blue) regions, and inclusion (bold) or skipping (dotted) of exonic or intronic sequences. (**D**) Predicted consequences of experimentally validated DSEs in *MAP3K7* or *CASP8*, highlighted correctly predicted NMD events in *MAP3K7* and domain alterations in *CASP8*. (**E**) Predicted consequences of experimentally validated DSE in *HRAS*, highlighted correctly predicted exon skipping (ES) events that leads to a longer oncogenic isoform in the truncated (ENST00000417302.5) and NMD (ENST00000493239.5) isoforms. A comparison of Isoform structures with the oncogenic isoform is provided in Supplementary Figure 2G. (**F**) Predicted consequences of *MPRIP* exon 23 skipping (ES) and inclusion (EI) in multiple Ref-TX show as % domain integrity (left). Domain integrity is defined as the percent of the original protein domain structure retained after the simulated event. ES simulation was conducted on ENST0000034712.8 and domain changes were identified in proteome and chain regions (right). Proteome (set of expressed proteins encoded by genome assembly) and chain (specific continuous regions of protein sequences) were defined by UniProt DB. (**G**) Predicted 3D protein structures of *MPRIP* based on the translated amino acid sequences of the Ref-TX (gray) and Sim-TX (brown), with amino acids encoded by the by the skipped exon 23 isoform highlighted in magenta and those from the included isoform in green.

Among the 118 NMD-sensitive Sim-TXs classified as ‘protein coding’ in GENCODE, 103 transcripts used a different reading frame from their Sim-TXs counterparts, while 15 transcripts were classified as ‘protein coding’ despite harboring a PTC in the same frame. We focused on the explainable 103 transcripts. For these, we applied the reading frame of each Sim-TX to its corresponding ‘protein coding’ transcripts (n=96, after filtering for availability) to determine whether a PTC was introduced within the coding sequence.

### Comparing protein structures

To analyze differences in 3D protein structure in *non-NMD* cases, we created a custom script ‘Make_aa_fa.py’ which extracts coding sequences from Ref-TX and Sim-TX FASTA files, and performs *in-silico* translation to generate amino acid sequence FASTA files. These amino acid sequence FASTA files are used to generate PDB files with AlpahFold2 (24). These resulting PDB structures are then aligned based on their shared amino acid sequences using PyMol (31) interactive mode, with differing amino acid residues highlighted with distinct colors.

### Comparing frequency of functional events

To compare the distribution of functional classes (LoD, GoD, NMD, and PTC removal) across FDR and PSI bins, we binned FDR (-log10 scale) and absolute *ΔPSI* values into 10 bins each. All rMATS FDR values of 0 were assigned a minimal FDR value of 2.2e-16 (the smallest FDR reported by rMATS). Within each bin, we counted the number of functional classes, and normalized these counts by the total number of DSEs per bin. In this analysis, we considered only the functional classes defined by the highest coding potential ORF (pORF1).

### Calculating an effect score

We defined an Effect Score to prioritize DSEs based on both functional impact and likelihood of occurrence:

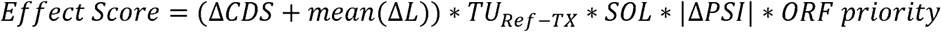

This score integrates five major components:

1. the functional consequence of the simulated splicing event, quantified by Δ*CDS* which represents the proportion of the coding sequence altered by the splicing event, estimating disruption of protein-coding regions, and Δ*L* which represents the mean change in length of annotated functional domain (*e.g.*, protein domains) between Ref-TX and Sim-TX.
2. the likelihood that the simulated event occurs in the given dataset, quantified as *transcript usage* (TU): *TU_i_* = *TPM_i_* / ∑*_kEG_ TPM_k_*, where *i* and *k* are transcripts of the same gene *G*. TU reflects the relative abundance and thus the likelihood that the event occurs in transcript *i*.
3. the splicing occurrence likelihood (*SOL*) is derived from the maximum PSI value from rMATS JCEC outputs. For inclusion cases, SOL is the higher PSI between groups; for skipping cases, its the lowest PSI.
4. the absolute PSI change (|Δ*PSI*|) measures the magnitude of the simulated splicing change between groups.
5. the ORF priority is assigned based on predicted coding potential, with scores of 1.0, 0.5, or 0.1 for the top three ranked pORFs, respectively.

Since all components are scaled between 0 and 1, the ‘Effect Score’ ranges from 0 to 2.

### Assigning the group with higher expression of Sim-TX

To facilitate interpretation of the functional consequences, we selected, for each DSE, the Ref-TX pair with the highest effect score as the representative event. For these, we assigned the group with higher Sim-TX expression based on the simulated splicing type (*i.e.,* skipping or inclusion) and *ΔPSI* direction:

1. for inclusion events (exon inclusion (EI), retained intron (RI), alternative A3/5SS (Alt_A3/5SS), and mutually exclusive exon1 (MXE1)) the group with higher mean PSI was considered to have higher Sim-TX expression.
2. for skipping event (exon skipping (ES), skipped intron (SI), canonical A3/5SS (Can_A3/5SS), and mutually exclusive exon2 (MXE2)), the group with lower mean PSI was considered to have higher Sim-TX expression.

### Gene ontology analysis

Gene ontology (GO) enrichment analysis were conducted using *Gseapy* (32) python module using GO_Biological_Process_2021 and Reactome_2022 terms. P-values were adjusted for multiple testing within *Gseapy*, and pathways with *P_adj_* <0.05 were considered significantly enriched.

Among the GO terms that were shared between SpliceDecoder and rMATS, we compared the -log_10_(P_adj_) values. Terms in which the SpliceDecoder value was at least 0.8 greater than the rMATS value were defined as SpliceDecoder-enriched terms, whereas terms in which the rMATS value was at least 0.2 greater than the SpliceDecoder value were defined as rMATS-only terms. The proportion of genes with functional changes within each term was defined by the fraction of DSEs exhibiting any functional consequences.

## Results

### SpliceDecoder workflow for differential splicing analysis

SpliceDecoder is designed to predict the functional consequences of differential splicing events (DSEs) identified using RNA-seq data processed through an event-based differential splicing analysis pipeline (**Fig. 1A**). In the first step, ‘Generate All Possible Splicing Cases’, the output of an event-based tools such as rMATS is used to infer two possible splicing outcomes—inclusion and skipping—for each splicing event type, including cassette exon (CA), alternative 5′ splice site (A5SS), alternative 3′ splice site (A3SS), retained intron (RI), and mutually exclusive exons (MXE) (**Fig. 1A** and **Supplementary Fig. 1A**). These inferred cases are used to construct a genomic coordinate matrix and identify target positions for finding matching isoforms in a given GTF file (**Supplementary Fig. 1A**, See **Methods**). In the second step, ‘Map Splicing Cases’, SpliceDecoder matches these splicing cases to transcripts in the GTF based on the identified target positions and designates these perfectly matched transcripts as Ref-TXs (**Supplementary Fig. 1B)**. For example, for exon skipping (ES) events in DSE#1, matched transcripts are labelled as ES-Ref-TX (**Fig. 1A**). In the third step, ‘Simulate Splicing Events’, a given splicing event is simulated within a full-length transcript using the structure of the matched Ref-TX as template. These include exon inclusion (EI), exon skipping (ES), retained intron (RI), skipped intron (SI), alternative 3’/5’ splice site (Alt_A3/5SS), canonical 3’/5’ splice site (Can_A3/5SS), and mutually exclusive exon 1/2 (MXE1/2). For instance, adding a previously skipped exon to a Ref-TX (ES-Ref-TX_1_) yields a simulated inclusion transcript (EI-Sim-TX_1_) (**Fig. 1A**). The fourth step, ‘Functional Annotation’, involves annotating functional features such as protein domains, regions, motifs, DNA binding sites, disordered regions and the potential for nonsense-mediated decay (NMD), using the genomic coordinates of Ref-TX and Sim-TX (See **Methods**). Functional impacts are then predicted by comparing the annotated features of Ref-TX and Sim-TX (**Fig. 1A**). Finally, the fifth step, ‘DSEs with Effect Score’, SpliceDecoder outputs 9 summary files. These include effect scores, visual representations of Ref-TX and Sim-TX, reports on domain alterations and NMD predictions, and the amino acid sequences of both Ref-TX and Sim-TX (**Fig. 1A**).

We applied SpliceDecoder to our test dataset which used previously described RNA-seq from human mammary epithelial MCF-10A cells at 0, 8, and 24 hours after MYC activation (n=3) (14). SpliceDecoder successfully mapped 9,124 (89%) and 8,025 (88%) DSEs from the corresponding rMATS outputs, respectively, to at least one Ref-TX, with on average, each DSE mapping to 4 Ref-TXs. (**Fig. 1B, C** and **Supplementary Fig. 1C**). Using these DSE and their corresponding Ref-TXs, we generated simulated transcripts (Sim-TX) containing alternative splicing events. For downstream functional analysis, we selected the top three open reading frames (ORFs) with the highest CPAT2-derived coding potential (CP) scores as default candidates (**Fig. 1D** and See **Methods**) (26).

### Predicting the functional consequences of DSEs based on simulated splicing

To predict the functional consequences of DSEs, we developed an algorithm that classifies them into one of seven functional classes: *Loss of Frame* (LoF), *NMD*, *PTC removal*, *Loss of Domain* (LoD), *Gain of Domain* (GoD), *CDS alteration* (CDS alt), and *UTR alteration* (UTR alt) (**Fig. 2A**). These classifications are assigned by comparing Ref-TX and Sim-TX for each DSE-Ref-TX pair. There are four key steps to achieve these functional annotations:

1. ‘Frame Check’ Step: to minimize translational variations from reading frame differences, we first assess whether Sim-TXs match the start and stop codons of Ref-TX based on genomic coordinates (See **Methods**). Sim-TXs with a perfectly or partial frame match are classified as *Intact Frame*, while those lacking a matched start codon and a matched stop codon are classified as *Loss of Frame* (LoF) (**Fig. 2A**).
2. ‘NMD Check’ Step: we investigate whether *Intact Frame* Sim-TX can be targets for NMD or remove a premature termination codon (PTC) existing in the Ref-TX (**Fig. 2A**). NMD targets are defined by the ‘50-55 rule’, where a PTC occurs more than 55 nucleotides (nt) upstream of the last exon-exon junction (29). Based on the position of stop codon and last exon-exon junction, we classify *Intact Frame* cases as *NMD*, *PTC removal*, or *non-NMD*. For example, in *SRSF3*, simulated exon inclusion (simEI chr6;36599821;36600276) introduces a PTC in Sim-TX, classifying it as *NMD* (**Fig. 2B**); whereas in *SRSF7*, simulated exon skipping (simES chr2;38749346;38748898) eliminates a PTC in the Ref-TX, classifying it as *PTC removal* (**Fig. 2C**). To explore additional cases potentially escaping NMD, we also applied an advanced criterion, where a PTC located less than 150 nt downstream of the start codon or within an exon longer than 407 nt are classified as *non-NMD* (30) (See **Methods**). As a result, 5,719 cases that were previously classified as *NMD* by the basic criterion were reclassified as *non-NMD* by this advanced criterion (**Supplementary Table 3**).
3. ‘Domain Check’ Step: for *non-NMD* cases, we analyzed annotated functional features - domains, regions, sites of post-translational modifications, UTR regions, motifs, or binding sites - using UniProt DB (27) (**Fig. 2A**). We then classify changes into *Loss of Domain* (LoD), *Gain of Domain* (GoD), or *Intact Domain* and report the corresponding domain alterations in the *Domain_Alts.tsv* file (**Fig. 1A** and **Supplementary Table 4**). To account for length bias, we define a functional change ratio (Δ*L*) for each feature (See **Methods**). For example, in *CHEK1*, the simulated exon skipping of (simES chr11;125637449; 125637544) maintains the canonical reading frame, but results in the loss of the protein kinase and the CLSPN interaction region (Δ*L*=0.037) (**Fig. 2D** and **Supplementary Table 4**). Conversely, in *IRF3*, simulated inclusion of a mutually exclusive exon (simMXE1 chr19;49664846;49664674) introduces a new start codon and additional domains (Δ*L*=0.57) (**Fig. 2E** and **Supplementary Table 4**). Finally, NMD events, like those in *SRSF3* and *SRSF7*, are assigned a default Δ*L* of 1.
4. ‘Coding Sequence Length Check’ Step: for *Intact Domain* cases, we investigate coding sequence differences (Δ*CDS*) between Ref-TX and Sim-TX. Cases with non-zero Δ*CDS* (Δ*CDS* > 0) are classified as *CDS alterations* (CDS alt), while others are classified as *UTR alterations* (UTR alt) (**Fig. 2A**).

To aid interpretation, SpliceDecoder provides graphical representations of Ref-TX and Sim-TX pair, highlighting domain differences (**Fig. 2B-E**). Additionally, a tab-separated values (TSV) table lists the used ORF, CDS differences, UTR differences, Δ*L* scores, and the functional class for each DSE and Ref-TX pair in the *Main_Table.tsv* (**Fig. 1A** and **Supplementary Table 5**). The full-length amino acid sequences of both Ref-TX and Sim-TX can also be used as input for structure prediction tools such as AlphaFold2, facilitating downstream investigation of DSE-induced structural changes (**Fig. 2D**, **E**).

In addition to event-centric results, SpliceDecoder also supports the interpretation of transcript-centric results by applying the same classification algorithm to investigate functional differences between automatically defined canonical transcripts and all other transcripts within the same gene (**Fig. 2A** and See **Methods**). Using this transcript comparison function, we compared transcript-level data of *SRSF2* and *SRSF7*, and produced well-curated functional differences for each query pair (**Supplementary Fig. 2A**, B).

### Distribution of predicted functional consequences

We examined the frequency of each functional class (*NMD*, *PTC removal*, *LoD*, *GoD*, *CDS alt*, *UTR alt*, *LoF*) for each splicing event type (CA, RI, A3SS, A5SS, and MXE) in our MYC 8h and 24h datasets (**Fig. 2F**). In total, we observed 8,971 *NMD* (27%), 1,365 P*TC removal* (4%), 4,603 *LoD* (14%), 4,753 *GoD* (14%), 3,497 *CDS alt* (11%), 9,701 *UTR alt* (29%) and 247 *LoF* (0.7%) events. Retained introns showed a significantly higher occurrence of *NMD* events (n=3,521, 55% of RI events), as expected since these events often introduce PTCs (n=33,34). Retained introns also exhibited fewer *UTR alt* (n=1,470) and *CDS alt* (n=422) events (n=1,892, 29% of RI events) compared to other splicing event types (**Fig. 2F** and **Supplementary Fig. 3A**). All splicing events showed *UTR alt* classes, ranging from 20 to 50% of all events, suggesting not all DSEs affect protein coding sequences (**Fig. 2F**). We reasoned that the high proportion of *UTR alt* events may result from multi-mapping of DSEs, where a single DSE corresponds to multiple Ref-TXs (**Fig. 1B**), not all of which cause functional changes. Indeed, we found that 49% of DSEs exhibit different functional consequences depending on their Ref-TX (**Supplementary Fig. 3B**) with 4,994 DSEs (70%) causing at least one functional change across their Ref-TXs (**Fig. 2G** and **Supplementary Fig. 3C**). This data highlights the complexity of DSE functional consequences and underscores the importance of accurate Ref-TX mapping in functional interpretation.

In addition to examining the distribution of functional classes, we investigated frequently altered functional features by DSEs using the *Domain_Alts.tsv* file. We found that 24,940 regions, 16,476 domains, 1,549 binding sites, and 379 motifs were affected by different DSEs (**Fig. 2H**). We further investigated the top 10 domains and motifs and observed that similar features were frequently altered across different splicing types (**Supplementary Fig. 3D**). For example, KRAB, protein kinase, RRM, and C2H2-type domains, as well as nuclear localization signal motifs were consistently ranked among the top in multiple splicing types. Notably, these domains are involved in RNA binding, splicing regulation, and transcriptional control; thus, alterations in these domains may impact transcriptional programs and cellular processes. These findings provide a novel approach for interpreting and understanding the impact of DSEs at the protein domain level within the given dataset.

Finally, we investigated the relationship between Ref-TX CDS length and DSE functional classes. Interestingly, *LoD* and *NMD* cases were significantly more prevalent in longer Ref-TX (**Fig. 2I** and **Supplementary Fig. 3E**); while *GoD* and *PTC removal* cases were more frequent in shorter Ref-TX (**Fig. 2I** and **Supplementary Fig. 3E**). These patterns resemble the CDS length distribution of canonical and non-canonical isoforms. These findings suggest that DSEs occurring on non-canonical transcripts are more likely to increase their functionality; and conversely, DSEs on canonical transcripts are more likely to decrease their functionality.

### Validating the predicted functional consequences of DSEs

To assess the functional predictions made by SpliceDecoder, we examined 2,542 NMD-inducing DSEs from the MYC 8h dataset. We identified 488 GENCODE transcripts with intron chains perfectly matching those of their corresponding NMD-sensitive Sim-TXs (**Fig. 3A**). Of these, 370 transcripts (76%) were classified by GENCODE as NMD-sensitive transcripts, including 219 NMD-transcripts and 151 retained intron transcripts (**Fig. 3B**). We further investigated the remaining 118 protein coding transcripts (24%) and found that the majority (n=103) used a different reading frame than their corresponding NMD-sensitive Sim-TXs (See **Methods**). To account for these differences, we realigned their reading frames with the corresponding Sim-TXs and performed *in silico* translation. This analysis showed that all protein-coding transcripts with available sequence data (n=96) contained a PTC (**Fig. 3B**). Consequently, these findings confirmed that 466 (95%) of the NMD-sensitive Sim-TXs predicted by SpliceDecoder were either annotated as NMD-sensitive or showed strong evidence of NMD targeting in GENCODE.

Next, we evaluated SpliceDecoder on two curated set of poison exons which are a class of exons known to introduce PTCs and reduce protein production (35). These included experimentally validated poison exons in 14 splicing factors from the SR protein family (36–39), and 1,583 computationally predicted poison exons in 1,198 RNA binding proteins (RBPs). From the MYC 8h dataset, we extracted corresponding DSEs at standard cut-offs (FDR<0.05 and |ΔPSI|>0.1) and compared their predicted functional consequences to those previously experimentally validated (35). Among the SR protein poison exons, 7 matched DSEs were identified in our dataset, and all were correctly classified by SpliceDecoder as *NMD* due to poison exon inclusion (**Supplementary Table 6**). These Sim-TXs were assigned the maximal Δ*L* score of 1.0, indicating complete loss of Ref-TX functional domains (**Fig. 3C** and **Supplementary Fig. 3F**). Furthermore, among the computationally predicted RBP poison exons, 113 matched DSEs were found in our MYC dataset. Of these, 81 (72%) were correctly classified as *NMD* or *PTC removal* events, such as the skipping of the *EIF4A2* poison exon which eliminates a PTC (**Supplementary Fig. 3G**). However, 32 (28%) were not classified as *NMD* or *PTC removal*, prompting further investigation. We found that these exons are predicted to induce NMD only in specific NMD-sensitive transcripts. Because event-based approaches like rMATS report DSEs without isoform context, SpliceDecoder may assigns a non-NMD-sensitive Ref-TX transcript that shares the same exon structure, thereby simulating the event in a non-NMD background and predicting it as a non-NMD outcome (**Supplementary Fig. 3H**). These differences result from the inherent limitations to event-centric approaches from splicing detection.

Lastly, we examined on a set of experimentally validated DSEs known to alter protein domain (15,40,41). We first analyzed two events: an alternative 3’ splice site in *MAP3K7* and an exon skipping in *CASP8* (40), both previously shown to trigger NMD and affect domain architecture (40). SpliceDecoder correctly predicted that the alternative 3’ splice site event in *MAP3K7* introduces a PTC within the middle of Ref-TX (ENST00000369325.7); while skipping of exons 6 and 7 of *CASP8* resulted in *LoD* in the caspase family p20 domain compared to the inclusion transcript (**Fig. 3D**). Next, we investigated an exon skipping in *HRAS* which produces a longer oncogenic RAS isoform (ENST00000311189.8) (41), that has been detected in breast and prostate tumors with high MYC activity (14,42). SpliceDecoder correctly predicted that exon skipping in both the NMD-sensitive (ENST00000493230.5) and truncated (ENST00000417302.5) transcripts would result in longer isoforms structurally matching the known oncogenic isoform (41) (ENST00000311189.8) (**Fig. 3E** and **Supplementary Fig. 3I**). Finally, we analyzed exon skipping in *MPRIP*, previously validated in metastatic pancreatic cancer (15), which alters phosphorylation sites and protein structure. SpliceDecoder predicted that exon skipping (ES) increases total domain length in two Ref-TXs, while exon inclusion (EI) reduces total domain length in five Ref-TXs (**Fig. 3F**), consistent with previous findings (15). At the individual domain level, exon skipping caused variations between residues S1016 and D1038 (**Fig. 3F**), again in agreement with published findings (15). To further investigate structural changes induced by *MPRIP* exon skipping, we used AlphaFold2 (24) to model the translated sequences of Ref-TX and Sim-TX. The analysis revealed an additional α-helical structure at the C-terminus of Sim-TX protein (**Fig. 3G**), consistent with structural differences previously reported (15).

In sum, SpliceDecoder correctly predicted functional consequences of alternative splicing across multiple datasets, demonstrating its utility in linking RNA-seq-based splicing changes to downstream impacts on protein structure and function.

### Developing a combinatorial score for measuring splicing and functional effects

Next, we evaluated whether conventional splicing measurements — specifically percent of spliced-in (PSI) and false discovery rate (FDR) — accurately capture the functional consequences of DSEs. To test this, we binned DSEs by FDR and ΔPSI (PSI of group2 – PSI of group1) and compared the number and frequency of functionally impactful DSEs, including *LoD*, *GoD*, *NMD* and *PTC removal* in each bin (**Fig. 4A, B**). Surprisingly, we observed that a higher number of functionally relevant DSEs occurred in bins with lower -log_10_FDR and ΔPSI values (**Fig. 4A, B** and **Supplementary Fig. 4A**, B). These patterns suggest that the most significant splicing events may not necessarily correspond to the most functionally impactful ones. Consequently, relying solely on FDR and ΔPSI to prioritize DSEs might lead to misleading conclusions.

**Figure 4.**
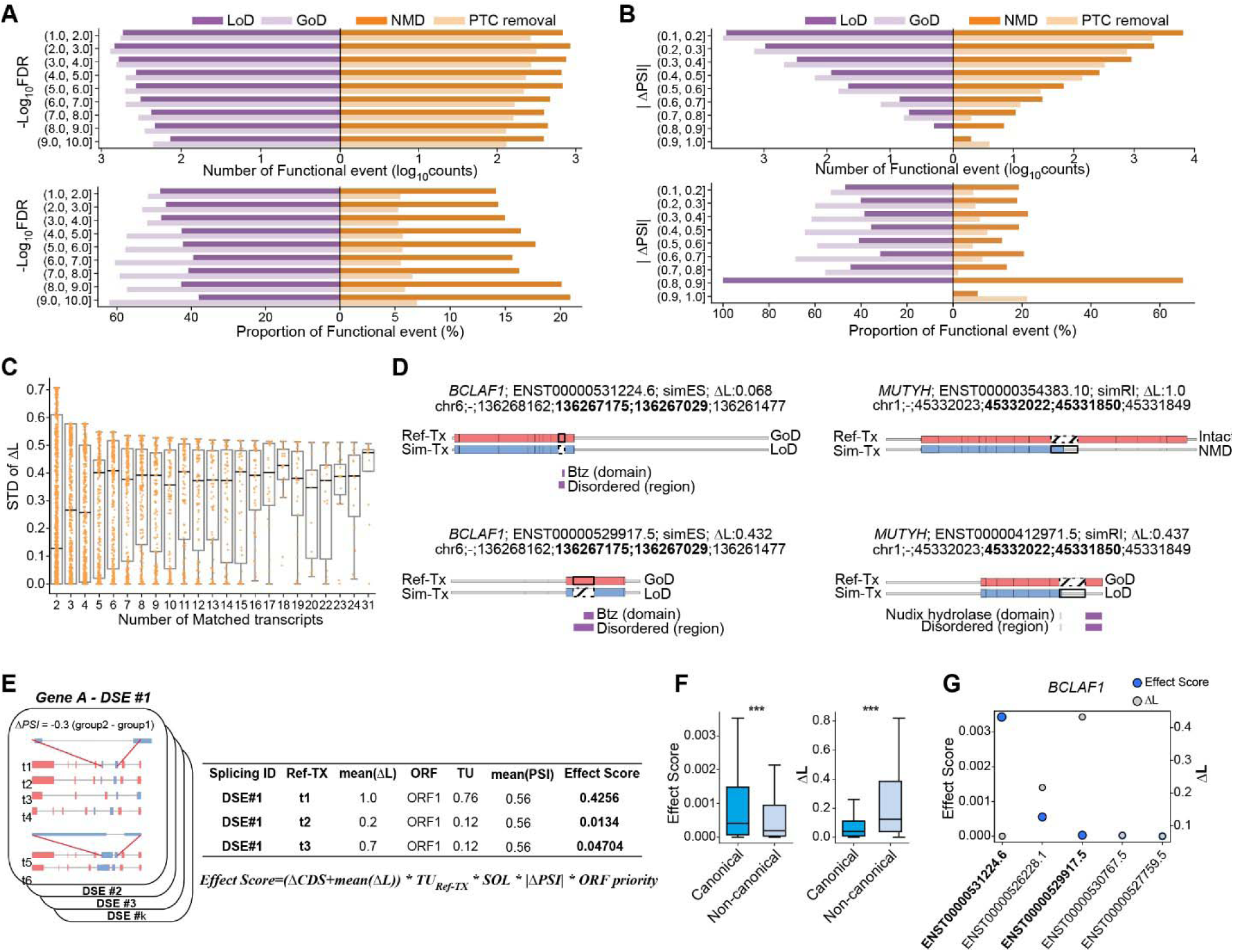
Association between the frequency of functional changes and conventional indices and flow chart of the combinatorial function process. (**A, B**) Association between conventional splicing analysis measurements such as FDR (**A)** or ⊿PSI (**B**) and the number (upper) and proportion (lower) of SpliceDecoder functional events, shown by functional classes (LoD, GoD, NMD, PTC removal), for our test dataset comparing human mammary epithelial MCF-10A cells at 8h after MYC activation vs. control. (**C**) Standard deviation (STD) of ⊿L for individual Ref-TX within same DSE. The x-axis indicates the number of matched transcripts for each DSE, and each dot represents a STD of individual DSE event. The bold black line represents the median value within each boxplot. (**D**) Example of different ⊿L consequences in the same DSE. In *BCLAF1*, simulated exon skipping (SimES) exhibits a lower ⊿L in the canonical isoform (ENST00000531224.6) but a higher ⊿L consequence in the non-canonical isoform (ENST00000529917.5). In *MUTYH*, simulated retained intron (SimRI) is predicted to lead to NMD in the canonical isoform (ENST00000354383.10) but cause LoD in the non-canonical isoform (ENST00000412971.5). (**E**) Schematic of the Effect score calculation process, which is defined as the combined formula of ΔCDS, mean ⊿L, transcript usage of Ref-TX (TU_Ref-TX_), splicing occurrence likelihood (SOL), ΔPSI, and ORF priority (See **Methods**). (**F**) Comparison of Effect scores and ⊿L between canonical and non-canonical isoforms in 221 DSEs, where the non-canonical isoforms have higher ⊿L. While non-canonical isoforms have significantly higher ⊿L, canonical isoforms exhibit significantly higher Effect score when accounting for various biological features. Statistical significance was estimated using Mann-Whitney U test. (**G**) Effect score and ⊿L for each DSE and Ref-TX pair for the *BCLAF1* gene. The Effect score identifies the canonical isoform (ENST00000531224.6) as the highest DSE and Ref-TX pair. Statistical significances were calculated using a one-sided Mann-Whitney U test.

To address this limitation, we examined whether the functional change ratio (*ΔL*) could serve as an alternative metric. However, when analyzing the standard deviation (STD) of *ΔL* across different Ref-TX pairs for the same DSE, we observed significant variation (**Fig. 4C**). This data indicates that a single DSE can produce significantly different functional outcomes depending on the Ref-TX context. For example, in *BCLAF1*, simulated exon skipping (simES) in the canonical isoform (ENST00000531224.6) resulted in a domain loss with minimal functional change (*ΔL*=0.068), whereas the same DSE in the non-canonical isoform (ENST00000529917.5) resulted in the same domain loss but exhibited a larger functional change due to differences in Ref-TX domain length (*ΔL*=0.432, **Fig. 4D**). Similarly, in *MUTYH*, a simulated retained intron (simRI) triggered NMD in ENST00000354383.10 (*ΔL*=1.0), but caused a modest domain loss in ENST00000412971.5 (*ΔL*=0.437, **Fig. 4D**). These findings highlight the importance of considering both the magnitude of *ΔL* and the specific Ref-TX context when interpreting functional consequences.

To improve prioritization, we developed a combined effect score that integrates: (1) the predicted functional consequence, and (2) the likelihood of the simulated splicing event (See **Methods**). This score enables the identification of the most biologically significant DSE and Ref-TX pair in each event (**Fig. 4E** and **Supplementary Table 7**). Notably, when considering the top 100 DSEs based on the effect score, most DSEs were annotated as NMD-related functional classes (*NMD* or *PTC-removal*), which are related to transcript degradation and influence gene expression level. In contrast, selecting the top 100 DSEs based on |ΔPSI| captured a higher frequency of *CDS alt* class, which may represent variability rather than functional impact (**Supplementary Fig. 4C**). We then tested the performance of the effect score by comparing canonical and non-canonical isoforms across 221 DSEs where non-canonical isoforms showed higher *ΔL* values (**Fig. 4F**). Despite their higher Δ*L*, non-canonical isoforms generally showed lower effect scores due to their lower expression or splicing likelihood, resulting in higher prioritization of canonical isoforms (**Fig. 4F**). For example, in *BCLAF1,* the non-canonical isoform (ENST00000529917.5) had a greater *ΔL*, but was expressed at very low levels, thus receiving a lower effect score compared to the canonical isoform (ENST00000531224.6) (**Fig. 4G**). Together, these findings demonstrate that the effect score prioritizes biologically meaningful DSE–Ref-TX pairs by combining functional impact with splicing context, and helps filter out misleading candidates that arise from relying on statistical or functional metrics alone.

### The effect score yields a more comprehensive view of the pathway impacted by alternative splicing

We next explored whether DSEs predicted by SpliceDecoder to have functional consequences are enriched in genes associated with specific biological pathways. Using differentially spliced gene lists from the MYC 8h and 24h datasets, we performed gene ontology analyses using Biological Process (43) and Reactome pathways (44). We compared two sets of DSEs: (1) those detected by rMATS using conventional splicing metrics alone (n=4,300 DSEs; FDR<0.05 and |ΔPSI|>0.1), versus (2) those predicted by SpliceDecoder to be functionally impactful (n=2,270 DSE; resulting in NMD or domain alterations, with FDR<0.05 and |ΔPSI|>0.1). Interestingly, SpliceDecoder-filtered DSEs were more significantly enriched in functional pathways than the conventional rMATS list (**Fig. 5A** and **Supplementary Fig. 5A**), but uniquely highlighted biological pathways that were absent from the conventional analysis, such as *Signaling by ERBB2*, *Signaling by FGFR1*/*FGFR2*, and *Signaling by hedgehog* (**Fig. 5B** and **Supplementary Fig. 5B**, D). Additionally, pathways uniquely enriched in the rMATS list had a lower proportion of functionally relevant DSEs, whereas those enriched in the SpliceDecoder list showed a higher proportion of predicted functional effects (**Fig. 5C** and **Supplementary Fig. 5C**-G). These data suggest that enrichment results from the conventional rMATS list are biased by the inclusion of >2,000 DSEs likely to not have a biological impact - leading to misleading pathways associations (*e.g.*, mRNA transport, nuclear export, and gene expression). In contrast, filtering out non-functional DSEs using SpliceDecoder improved pathway specificity and biological interpretability.

**Figure 5.**
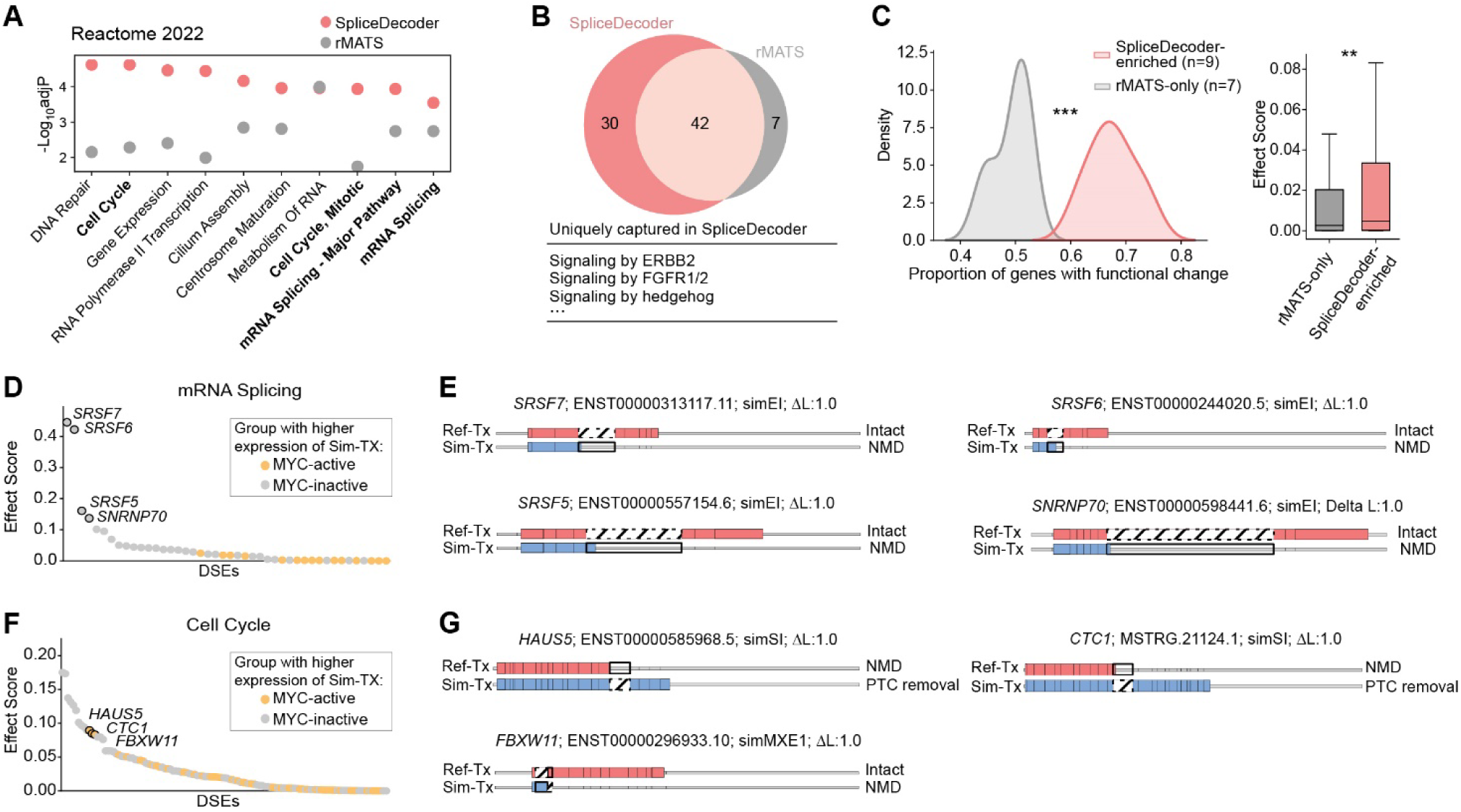
Difference in gene ontology analysis between rMATS and SpliceDecoder DSE gene lists. (**A**) Top 10 shared terms from a gene set enrichment analysis using either the spliced genes from a conventional rMATS DSE list (n=4,300; FDR<0.05, |ΔPSI|>0.1) or from the SpliceDecoder DSE list (n=2,270; NMD and domain alterations) for our test dataset comparing human mammary epithelial MCF-10A cells at 8h after MYC activation vs. control. (**B**) Overlap in enriched Reactome terms from the SpliceDecoder vs. rMATS spliced genes from **(A)**, with the top unique terms from SpliceDecoder shown below. (**C**) Proportion of genes with functional change and effect scores of ‘SpliceDecoder enriched’ or ‘rMATS only’ terms. ‘SpliceDecoder enriched’ includes terms with greater significance in SpliceDecoder enrichment compared to rMATS and ‘rMATS only’ includes terms uniquely significant in rMATS enrichment test. Statistical significance was estimated by using 2 sample KS test and one-sided Mann-Whitney U test. (**D, E**) Effect score for DSE and Ref-TX pairs impacting genes associated with ‘mRNA splicing’ term in the SpliceDecoder DSE list in MCF-10A cells at 8h after MYC activation vs. control (**D**). Examples of DSE and Ref-TX pairs with an effect score > 0.1 impacting genes associated with ‘mRNA splicing’ term and their predicted consequences (**E**). Gene symbol, Ref-TX ID, simulated event and ⊿L is shown for each event along with the transcript and exon structures of Ref-TX and Sim-TX with UTRs (gray), CDS (pink and blue) regions, and inclusion (bold) or skipping (dotted) of exonic or intronic sequences. (**F, G**) Effect score for DSE and Ref-TX pairs impacting genes associated with ‘Cell Cycle’ term in the SpliceDecoder DSE list in MCF-10A cells at 8h after MYC activation vs. control (**F**). Examples of DSE and Ref-TX pairs with an effect score > 0.08 impacting genes associated with ‘Cell Cycle’ term and their predicted consequences (**G**). Gene symbol, Ref-TX ID, simulated event and ⊿L is shown for each event along with the transcript and exon structures of Ref-TX and Sim-TX with UTRs (gray), CDS (pink and blue) regions, and inclusion (bold) or skipping (dotted) of exonic or intronic sequences.

We further focused on DSEs associated with the *mRNA Splicing* and *Cell Cycle* pathways – two of the most frequently represented *Reactome* categories (**Fig. 5A**). To clarify the functional consequences of individual DSE, we identified a representative DSE-Ref-TX pair for each event based on the highest effect score and assigned the group (MYC active or inactive) with higher Sim-TX expression (See **Methods** and **Supplementary Fig. 5H**). Within the *mRNA Splicing* pathway, several splicing factors, including *SRSF7*, *SRSF6*, *SRSF5*, and *SNRPN70*, had DSEs predicted to show increased poison exon inclusion in MYC-inactive cells (**Fig. 5D**). In all four cases, inclusion of the poison exon occurred within the canonical isoform (*SRSF7*-ENST00000313117.11, *SRSF6*-ENST00000244020.5, *SRSF5*-ENST00000557154.6, and *SNRNP70*-ENST00000598441.6) and SpliceDecoder predicted that the resulting included Sim-TXs would trigger NMD (**Fig. 5E**). These data suggest that poison exon inclusion in MYC-inactive cells promotes degradation of splicing factor transcripts via NMD, providing a potential mechanism for the elevated expression of splicing factors observed in MYC-active cancers. In contrast, DSEs the *Cell Cycle* pathway, such as those in *HAUS5*, *CTC1,* or *FBXW11*, were predicted to be more highly expressed in MYC-active cells (**Fig. 5F**). These DSEs either removed or introduced a PTC (*e.g.*, skipped introns or mutually exclusive exons), resulting in a switch between NMD-sensitive and protein coding isoforms (**Fig. 5G**). These data suggest that alternative splicing in MYC-active cells may promote the production of functional protein-coding isoforms of cell cycle related genes, potentially enhancing proliferative capacity.

In sum, the SpliceDecoder effect score — by integrating conventional splicing metrics with predicted functional impact — enables more accurate functional annotation of DSEs. This combined approach filters out non-functional events, sharpens pathway enrichment analyses, and helps prioritize splicing changes with the greatest biological relevance.

## Discussion

Here, we developed SpliceDecoder (https://github.com/hyeon9/SpliceDecoder/), a high-throughput splicing interpreter, that uses genomic coordinates to simulated alternative transcript isoforms for each DSE and prioritize those the most likely to have functional consequences. By identifying the best-matching simulated transcript (Sim-TX) for each DSE, SpliceDecoder predicts functional changes by integrating high-quality annotations of protein domains, DNA/RNA binding, linker regions, coding sequences and NMD likelihood. Based on these data, we identified new insights into how transcript length relates to splicing functional outcomes. Additionally, SpliceDecoder can also be applied to transcript-centric results, using the same classification algorithm to provide interpretation for the given full-length isoforms. Finally, by enabling integration with structural prediction tools like AlphaFold2, SpliceDecoder links RNA splicing variation to predicted protein structural changes. To further prioritize high-impact DSEs, we develop an effect score for each DSE and transcript pair, which incorporates both the predicted functional consequences, the magnitude and the reproducibility of splicing changes. We show that the effect score filters out likely non-functional DSEs, and enhances the identification of biologically relevant pathways impacted by alternative splicing. For broader usability, SpliceDecoder has been tested for compatibility with well-established RNA-seq pipelines (23), and efforts are underway to ensure support for other event-centric approach, such as SUPPA2 (45).

Despite its strengths, SpliceDecoder has some limitations. First, while the current effect score provides a useful prioritization strategy, it may not fully capture rare or complex functional cases. To accommodate this, SpliceDecoder outputs a complete set of predicted functional annotations, allowing users to apply custom ranking criteria tailored to specific biological contexts. Second, SpliceDecoder relies available transcript annotations (GTF) and known protein domain databases. However, it supports easy integration of user-defined annotations, including long-read-based isoform data, which are expected to improve in accuracy and availability with ongoing advances in long-read sequencing technologies. Finally, although SpliceDecoder was initially designed for short-read RNA-seq—which remains the most widely used approach for splicing analysis—its framework can be adapted for use with long-read datasets in the future.

In sum, SpliceDecoder is a versatile and powerful tool for prioritizing DSEs from short-read RNA-sequencing data and nominating those most likely to result in biologically meaningful functional consequences, helping to address a critical bottle neck in RNA splicing analysis. Moreover, its compatibility with transcript-centric results extends its utility to long-read RNA-sequencing platforms, offering a unified framework for transcriptome-wide interpretation of splicing alterations and their potential functional impact.

## Data Availability

The data underlying this article are available in the article and in its online supplementary material. The official documentation and source codes of the SpliceDecoder are available at https://github.com/hyeon9/Splice-decoder

## Supplementary data

Supplementary Data are available at NAR online.

## Acknowledgements

We thank members of the Anczukow and Chuang labs for helpful discussions. We acknowledge the use of high-performance computing resources provided by The Jackson Laboratory (JAX) and supported by the JAX Cancer Center (P30 CA034196).

## Author contributions

H.G.K., O.A. conceived and designed the project, performed the theoretical analyses, interpreted the data and wrote the manuscript. J.C. supervised the project and contributed to manuscript editing. M.Y. contributed to the development of the scoring function and the implementation of SpliceDecoder. M.B. provided feedback on the GENCODE-based validation analysis and the manuscript. H.G.K and O.A. wrote and edited the manuscript with input from all authors. All authors read and approved the final manuscript.

## Funding

This work was supported by National Institutes of Health (NIH) grants R00CA178206, R01CA248317 and R01GM138541 to OA. We acknowledge the use of shared resources supported by the JAX Cancer Center (NCI P30CA034196). The content is solely the responsibility of the authors and does not necessarily represent NIH official views.

## Conflict of interest

The authors declare no conflict of interest.

## Notes

### Competing Interest Statement

The authors have declared no competing interest.

## References

1. Wright, C.J., Smith, C.W.J. and Jiggins, C.D. (2022) Alternative splicing as a source of phenotypic diversity. Nat Rev Genet, 23, 697–710.

2. Baralle, F.E. and Giudice, J. (2017) Alternative splicing as a regulator of development and tissue identity. Nat Rev Mol Cell Biol, 18, 437–451.

3. Mazin, P.V., Khaitovich, P., Cardoso-Moreira, M. and Kaessmann, H. (2021) Alternative splicing during mammalian organ development. Nature Genetics, 53, 925–934.

4. Angarola, B.L. and Anczukow, O. (2021) Splicing alterations in healthy aging and disease. Wiley Interdiscip Rev RNA, 12, e1643.

5. Bradley, R.K. and Anczukow, O. (2023) RNA splicing dysregulation and the hallmarks of cancer. Nat Rev Cancer, 23, 135–155.

6. Nikom, D. and Zheng, S. (2023) Alternative splicing in neurodegenerative disease and the promise of RNA therapies. Nat Rev Neurosci, 24, 457–473.

7. Christofk, H.R., Vander Heiden, M.G., Harris, M.H., Ramanathan, A., Gerszten, R.E., Wei, R., Fleming, M.D., Schreiber, S.L. and Cantley, L.C. (2008) The M2 splice isoform of pyruvate kinase is important for cancer metabolism and tumour growth. Nature, 452, 230–233.

8. Israelsen, W.J., Dayton, T.L., Davidson, S.M., Fiske, B.P., Hosios, A.M., Bellinger, G., Li, J., Yu, Y., Sasaki, M., Horner, J.W. et al. (2013) PKM2 isoform-specific deletion reveals a differential requirement for pyruvate kinase in tumor cells. Cell, 155, 397–409.

9. Boise, L.H., Gonzalez-Garcia, M., Postema, C.E., Ding, L., Lindsten, T., Turka, L.A., Mao, X., Nunez, G. and Thompson, C.B. (1993) bcl-x, a bcl-2-related gene that functions as a dominant regulator of apoptotic cell death. Cell, 74, 597–608.

10. Dou, Z., Zhao, D., Chen, X., Xu, C., Jin, X., Zhang, X., Wang, Y., Xie, X., Li, Q., Di, C. et al. (2021) Aberrant Bcl-x splicing in cancer: from molecular mechanism to therapeutic modulation. J Exp Clin Cancer Res, 40, 194.

11. Shkreta, L., Michelle, L., Toutant, J., Tremblay, M.L. and Chabot, B. (2011) The DNA damage response pathway regulates the alternative splicing of the apoptotic mediator Bcl-x. J Biol Chem, 286, 331–340.

12. Guil, S., de La Iglesia, N., Fernandez-Larrea, J., Cifuentes, D., Ferrer, J.C., Guinovart, J.J. and Bach-Elias, M. (2003) Alternative splicing of the human proto-oncogene c-H-ras renders a new Ras family protein that trafficks to cytoplasm and nucleus. Cancer Res, 63, 5178–5187.

13. Huang, M.Y. and Cohen, J.B. (1997) The alternative H-ras protein p19 displays properties of a negative regulator of p21Ras. Oncol Res, 9, 611–621.

14. Urbanski, L., Brugiolo, M., Park, S., Angarola, B.L., Leclair, N.K., Yurieva, M., Palmer, P., Sahu, S.K. and Anczukow, O. (2022) MYC regulates a pan-cancer network of co-expressed oncogenic splicing factors. Cell Rep, 41, 111704.

15. Jbara, A., Lin, K.T., Stossel, C., Siegfried, Z., Shqerat, H., Amar-Schwartz, A., Elyada, E., Mogilevsky, M., Raitses-Gurevich, M., Johnson, J.L. et al. (2023) RBFOX2 modulates a metastatic signature of alternative splicing in pancreatic cancer. Nature, 617, 147–153.

16. Ma, W.K., Voss, D.M., Scharner, J., Costa, A.S.H., Lin, K.T., Jeon, H.Y., Wilkinson, J.E., Jackson, M., Rigo, F., Bennett, C.F. et al. (2022) ASO-Based PKM Splice-Switching Therapy Inhibits Hepatocellular Carcinoma Growth. Cancer Res, 82, 900–915.

17. Shin, M.S., Kirklin, J.K., Cain, J.B. and Ho, K.J. (1987) Primary angiosarcoma of the heart: CT characteristics. AJR Am J Roentgenol, 148, 267–268.

18. Wang, Z., Jeon, H.Y., Rigo, F., Bennett, C.F. and Krainer, A.R. (2012) Manipulation of PK-M mutually exclusive alternative splicing by antisense oligonucleotides. Open Biol, 2, 120133.

19. Zhang, J., Wang, Y., Li, S.Q., Fang, L., Wang, X.Z., Li, J., Zhang, H.B., Huang, B., Xu, Y.M., Yang, W.M. et al. (2020) Correction of Bcl-x splicing improves responses to imatinib in chronic myeloid leukaemia cells and mouse models. Br J Haematol, 189, 1141–1150.

20. Mehmood, A., Laiho, A., Venäläinen, M.S., McGlinchey, A.J., Wang, N. and Elo, L.L. (2020) Systematic evaluation of differential splicing tools for RNA-seq studies. Briefings in Bioinformatics, 21, 2052–2065.

21. Shen, S., Park, J.W., Lu, Z.-x., Lin, L., Henry, M.D., Wu, Y.N., Zhou, Q. and Xing, Y. (2014) rMATS: Robust and flexible detection of differential alternative splicing from replicate RNA-Seq data. Proceedings of the National Academy of Sciences, 111, E5593–E5601.

22. Pertea, M., Pertea, G.M., Antonescu, C.M., Chang, T.-C., Mendell, J.T. and Salzberg, S.L. (2015) StringTie enables improved reconstruction of a transcriptome from RNA-seq reads. Nature Biotechnology, 33, 290–295.

23. Ewels, P.A., Peltzer, A., Fillinger, S., Patel, H., Alneberg, J., Wilm, A., Garcia, M.U., Di Tommaso, P. and Nahnsen, S. (2020) The nf-core framework for community-curated bioinformatics pipelines. Nature Biotechnology, 38, 276–278.

24. Jumper, J., Evans, R., Pritzel, A., Green, T., Figurnov, M., Ronneberger, O., Tunyasuvunakool, K., Bates, R., Žídek, A., Potapenko, A., et al. (2021) Highly accurate protein structure prediction with AlphaFold. Nature, 596, 583–589.

25. Dobin, A., Davis, C.A., Schlesinger, F., Drenkow, J., Zaleski, C., Jha, S., Batut, P., Chaisson, M. and Gingeras, T.R. (2013) STAR: ultrafast universal RNA-seq aligner. Bioinformatics, 29, 15–21.

26. Wang, L., Park, H.J., Dasari, S., Wang, S., Kocher, J.P. and Li, W. (2013) CPAT: Coding-potential assessment tool using an alignment-free logistic regression model. Nucleic Acids Research, 41.

27. Bateman, A., Martin, M.J., Orchard, S., Magrane, M., Agivetova, R., Ahmad, S., Alpi, E., Bowler-Barnett, E.H., Britto, R., Bursteinas, B. et al. (2021) UniProt: the universal protein knowledgebase in 2021. Nucleic Acids Research, 49, D480–D489.

28. Quinlan, A.R. and Hall, I.M. (2010) BEDTools: a flexible suite of utilities for comparing genomic features. Bioinformatics, 26, 841–842.

29. Nagy, E. and Maquat, L.E. (1998) A rule for termination-codon position within intron-containing genes: when nonsense affects RNA abundance. Trends in Biochemical Sciences, 23, 198–199.

30. Lindeboom, R.G.H., Supek, F. and Lehner, B. (2016) The rules and impact of nonsense-mediated mRNA decay in human cancers. Nature Genetics, 48, 1112–1118.

31. Schrodinger, LLC. (2015).

32. Fang, Z., Liu, X. and Peltz, G. (2023) GSEApy: a comprehensive package for performing gene set enrichment analysis in Python. Bioinformatics, 39, btac757.

33. Baek, D. and Green, P. (2005) Sequence conservation, relative isoform frequencies, and nonsense-mediated decay in evolutionarily conserved alternative splicing. Proceedings of the National Academy of Sciences, 102, 12813–12818.

34. Middleton, R., Gao, D., Thomas, A., Singh, B., Au, A., Wong, J.J.L., Bomane, A., Cosson, B., Eyras, E., Rasko, J.E.J. et al. (2017) IRFinder: assessing the impact of intron retention on mammalian gene expression. Genome Biology, 18, 51.

35. Leclair, N.K., Brugiolo, M., Urbanski, L., Lawson, S.C., Thakar, K., Yurieva, M., George, J., Hinson, J.T., Cheng, A., Graveley, B.R. et al. (2020) Poison Exon Splicing Regulates a Coordinated Network of SR Protein Expression during Differentiation and Tumorigenesis. Mol Cell, 80, 648–665 e649.

36. Lareau, L.F., Inada, M., Green, R.E., Wengrod, J.C. and Brenner, S.E. (2007) Unproductive splicing of SR genes associated with highly conserved and ultraconserved DNA elements. Nature, 446, 926–929.

37. Best, A., Dagliesh, C., Ehrmann, I., Kheirollahi-Kouhestani, M., Tyson-Capper, A. and Elliott, D.J. (2013) Expression of Tra2 β in cancer cells as a potential contributory factor to neoplasia and metastasis. International Journal of Cell Biology.

38. Sureau, A., Gattoni, R., Dooghe, Y., Stévenin, J. and Soret, J. (2001) SC35 autoregulates its expression by promoting splicing events that destabilize its mRNAs. EMBO Journal, 20, 1785–1796.

39. Stoilov, P., Dauod, R., Nayler, O. and Stamm, S. (2004) Human tra2-beta1 autoregulates its protein concentration by infuencing alternative splicing of its pre-mRNA. Human Molecular Genetics, 13, 509–524.

40. Lee, S.C.-W., North, K., Kim, E., Jang, E., Obeng, E., Lu, S.X., Liu, B., Inoue, D., Yoshimi, A., Ki, M. et al. (2018) Synthetic Lethal and Convergent Biological Effects of Cancer-Associated Spliceosomal Gene Mutations. Cancer Cell, 34, 225–241.e228.

41. García-Cruz, R., Camats, M., Calin, G.A., Liu, C.-G., Volinia, S., Taccioli, C., Croce, C.M. and Bach-Elias, M. (2015) The role of p19 and p21 H-Ras proteins and mutants in miRNA expression in cancer and a Costello syndrome cell model. BMC Medical Genetics, 16, 46.

42. Phillips, J.W., Pan, Y., Tsai, B.L., Xie, Z., Demirdjian, L., Xiao, W., Yang, H.T., Zhang, Y., Lin, C.H., Cheng, D. et al. (2020) Pathway-guided analysis identifies Myc-dependent alternative pre-mRNA splicing in aggressive prostate cancers. Proceedings of the National Academy of Sciences, 117, 5269–5279.

43. Ashburner, M., Ball, C.A., Blake, J.A., Botstein, D., Butler, H., Cherry, J.M., Davis, A.P., Dolinski, K., Dwight, S.S., Eppig, J.T. et al. (2000) Gene Ontology: tool for the unification of biology. Nature Genetics, 25, 25–29.

44. Milacic, M., Beavers, D., Conley, P., Gong, C., Gillespie, M., Griss, J., Haw, R., Jassal, B., Matthews, L., May, B. et al. (2024) The Reactome Pathway Knowledgebase 2024. Nucleic Acids Research, 52, D672–D678.

45. Trincado, J.L., Entizne, J.C., Hysenaj, G., Singh, B., Skalic, M., Elliott, D.J. and Eyras, E. (2018) SUPPA2: Fast, accurate, and uncertainty-aware differential splicing analysis across multiple conditions. Genome Biology, 19.

