## Supplementary Figures for "SpliceDecoder: A High-Throughput Tool for Guiding the Functional Interpretation of Differential Splicing Events"

### **Supplementary Figure 1**. Overview of transcript matching process and mapping accuracy.

### **Supplementary Figure 2**. Functional comparisons across user-defined transcript pairs.

### **Supplementary Figure 3**. Biological clues for the estimated functional consequences in the additional dataset.

### **Supplementary Figure 4**. Association between the frequency of functional changes and conventional indices in the MYC 24h test dataset.

### **Supplementary Figure 5**. Different enriched pathways in several biological process.

**Supplementary Tables**

**Supplementary Table 1**. Example of the rMATS output of MYC 0h *vs.* 8h

**Supplementary Table 2**. Example of the rMATS output of MYC 0h *vs.* 24h

**Supplementary Table 3**. SpliceDecoder ‘NMD.tsv’ with advanced NMD criterion of MYC 0h vs 8h and 0h vs 24h

**Supplementary Table 4**. Example of the ‘Domain_Alts.txt’ for *CHEK1* and *IRF3*

**Supplementary Table 5**. SpliceDecoder ‘Main_Table.tsv’ of MYC 0h *vs* 8h and 0h *vs* 24h

**Supplementary Table 6**. Experimentally validated poison exons in splicing factors and their predicted functional consequences.

**Supplementary Table 7**. SpliceDecoder ‘effect score.tsv’ of MYC 0h *vs* 8h and 0h *vs* 24h

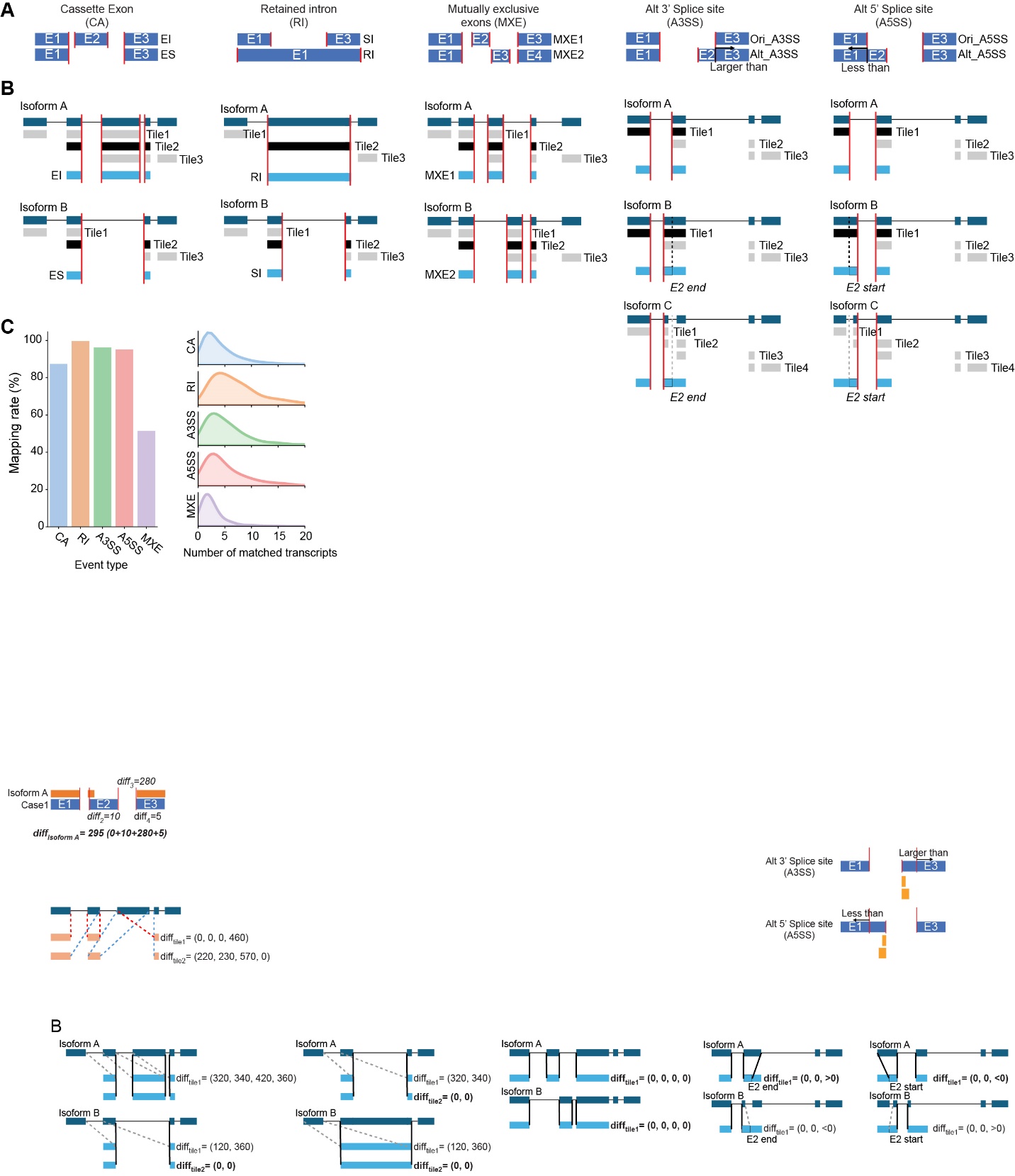

### **Supplementary Figure 1. Overview of transcript matching process and mapping accuracy.**

(**A**) Schematic representation of the ‘Target positions’ step for all possible splicing cases. Blue blocks represent exon structures for each splicing cases, red lines highlight target positions.

(**B**) Schematic representation of the ‘Map Splicing Case’ step for all possible splicing cases. Dark blue blocks represent exon structures for given isoforms (A and B), light blue blocks represent the given splicing cases, red lines represent the defined target positions, and black and gray blocks represent matched and unmatched exon structure tiles of each isoform. The dotted lines for A3/5SS categories indicate additional mapping criterion and the color of dotted lines indicate that satisfied case (black), unmet case (gray), respectively.

(**C**) Mapping rates of DSEs (left) and number of matched transcripts for query cases (right) shown per splicing event type for our test dataset comparing human mammary epithelial MCF-10A cells at 24h after activation of the MYC oncogene to a control cell line.

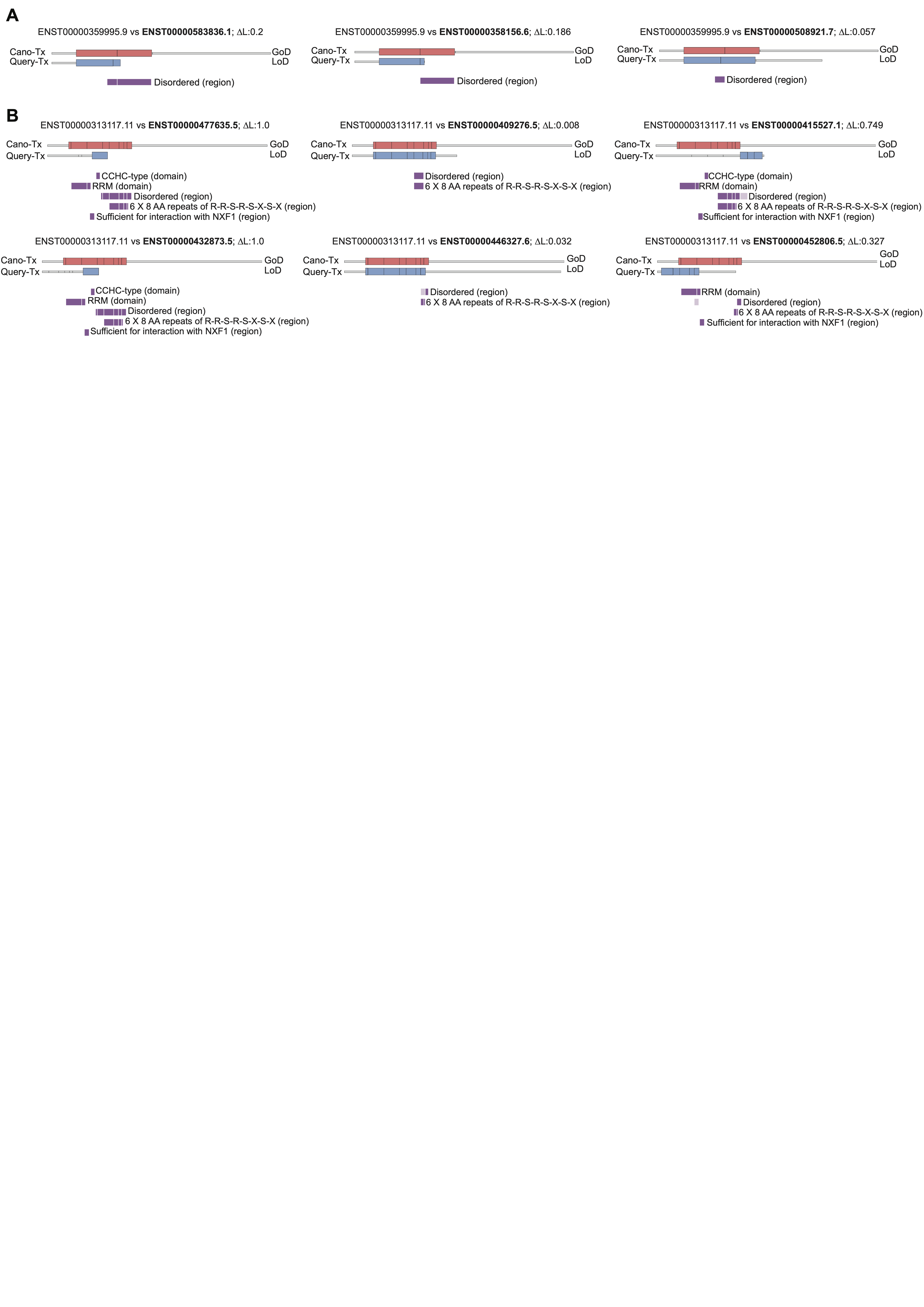

### **Supplementary Figure 2. Functional comparisons across user-defined transcript pairs.**

(**A, B**) Functional comparison of *SRSF2* (**A**) and *SRSF7* (**B**). For each comparison, the user-defined canonical transcript (Cano-Tx) ID, query transcript (Query-Tx, bolded) ID, and ∆L are shown above, with transcript and exon structures of the canonical and query transcript displayed below, with UTRs (gray) and CDS (pink and blue) regions. Lost domains in the query transcript are highlighted in dark purple, while gained domains are highlighted in light purple. Only comparisons with domain alterations are shown.

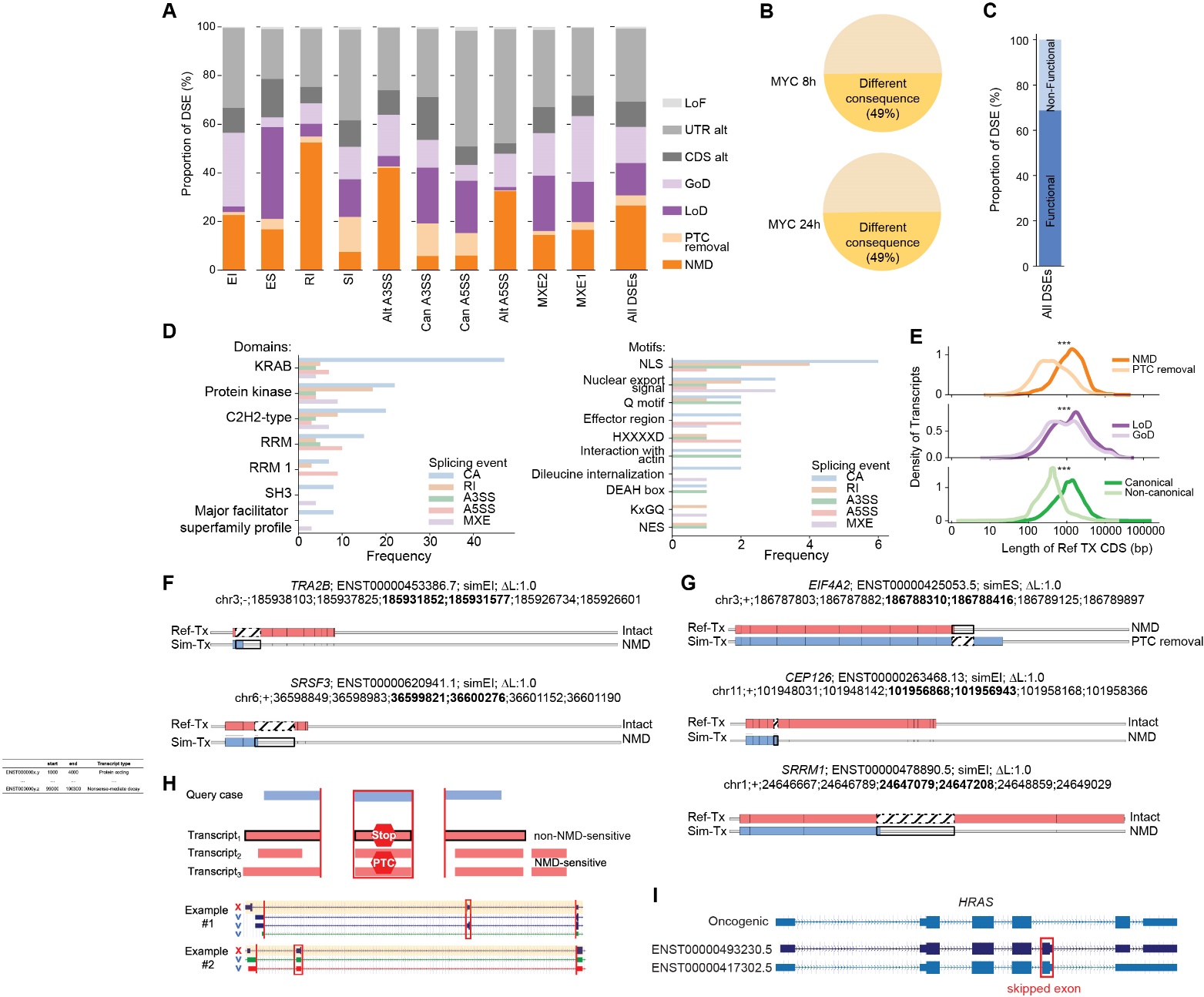

### **Supplementary Figure 3. Biological clues for the estimated functional consequences in the additional dataset.**

### (**A**) Distribution of each functional class per splicing event type (left) and for all events (right) for DSE detected in human mammary epithelial MCF-10A cells at 24h after MYC activation vs. control.

### (**B**) Proportion of DSEs with a different predicted consequence that are driven by difference in the selected Ref-TX in the MYC 8h and 24h test dataset.

### (**C**) Proportion of DSEs with at least one functional consequence (including LoD, GoD, NMD, and PTC removal) in human mammary epithelial MCF-10A cells at 8h or 24h after MYC activation vs. control.

### (**D**) Distribution of frequently altered domains (left) and motifs (right) across different splicing event types. Only domains and motifs altered by two or more different splicing types are shown.

### (**E**) Length of the matched transcript (Ref-TX) shown by functional consequences, NMD-related, PTC removal, LoD, and GoD, all classes were depicted as in **(A)**, and isoform types (Gencode canonical and non-canonical). Statistical differences were calculated by 2sample KS test.

### (**F**) Example of experimentally validated poison exons within splicing factors for which simulated exon inclusions were correctly classified as NMD.

### (**G**) Example of computationally predicted poison exon in RNA binding proteins. In *EIF4A2*, skipping of a poison exon (simES) is predicted to lead to PTC removal in the Sim-TX. In *CEP126* and *SRRM1*, inclusion of a poison exon (simEI) is predicted as an NMD class in Sim-TX.

### (**H**) Example of a poison exon that was not accurately classified as NMD because of differences in flanking exon structures. Blue blocks and red lines represent the exon structure of query case and the target positions respectively. The pink boxes with black outlines represent Ref-TX, while the other pink boxes represent unmatched transcripts. ‘Stop’ and ‘PTC’ labels indicate NMD-insensitive and NMD-sensitive isoforms respectively. In the two examples, the poison exon is incorrectly mapped only to the Ref-TX (marked with blue ‘v’) but not to the actual NMD-sensitive transcript (marked with red ‘x), and thus is not predicted as an NMD target.

### (**I**) Structure of the *HRAS* transcripts, with the red box indicating the skipped exon in **Fig. 3E,** which leads to expression of a more oncogenic isoform compared to Ref-TXs (ENST493230.5 and ENST417302.5).

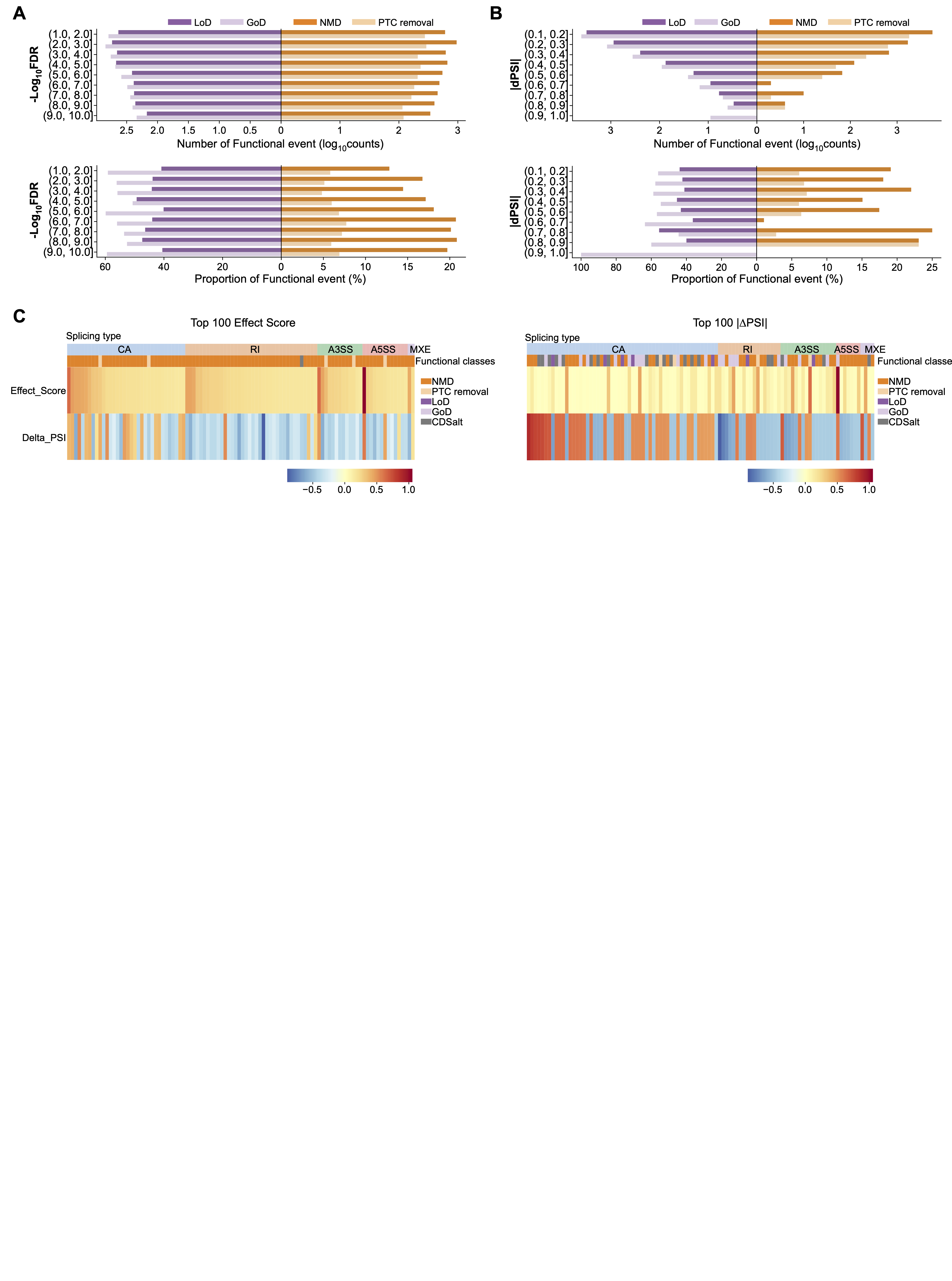

### **Supplementary Figure 4. Association between the frequency of functional changes and conventional indices in the MYC 24h dataset.**

(**A, B**) Association between conventional splicing analysis measurements FDR (A) or ⊿PSI ( B) and the number (upper) or proportion (lower) of SpliceDecoder predicted functional classes (LoD, GoD, NMD, PTC removal), for our test dataset comparing human mammary epithelial MCF-10A cells at 24h after MYC activation vs. control.

(**C**) Distribution of effect scores and *∆PSI* for the top 100 DSEs, with the top 100 DSEs ranked by effect score (left) or by *∆PSI* (right).

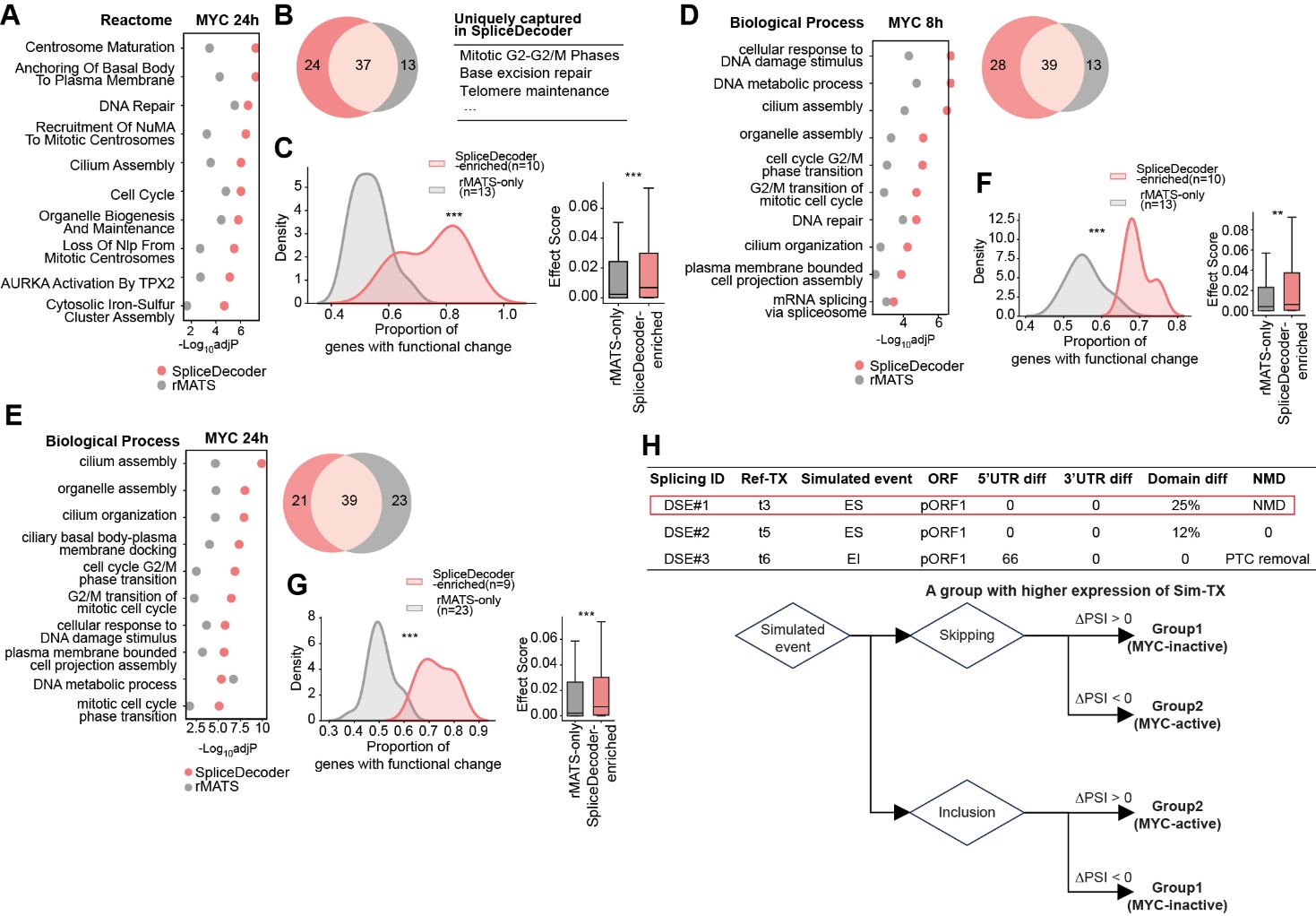

### **Supplementary Figure 5. Different enriched pathways in several biological processes.**

(**A**) Top 10 shared terms from a gene set enrichment analysis using either the spliced genes from a conventional rMATS DSE list (n=3,970; FDR < 0.05, |∆PSI| > 0.1) or from the SpliceDecoder DSE list (n=1,971; NMD and domain alterations) based on the Reactome 2022 terms for our test dataset comparing human mammary epithelial MCF-10A cells at 24h after activation of the MYC oncogene to a control cell line.

(**B**) Overlap in enriched Reactome terms of SpliceDecoder and rMATS from **(A),** highlighting the uniquely enriched terms in SpliceDecoder.

(**C**) Proportion of genes with functional change and effect scores of ‘SpliceDecoder-enriched’ and ‘rMATS only’ terms. ‘SpliceDecoder-enriched’ includes terms with greater significance in SpliceDecoder enrichment compared to rMATS and ‘rMATS only’ includes terms uniquely significant in rMATS enrichment test. Using a conventional rMATS DSE list (n=3,970; FDR<0.05, |ΔPSI|>0.1) for our test dataset comparing human mammary epithelial MCF-10A cells at 24h after MYC activation vs. control. Statistical significance was estimated by using 2sample KS test and one-sided Mann-Whitney U test.

(**D, F**) Gene set enrichment analysis results of rMATS and SpliceDecoder based on the GO_Biological_Process 2021 terms for MYC 8h or 24h vs. control, showing all shared terms between rMATS and SpliceDecoder. Venn-diagram represents shared and uniquely captured terms between SpliceDecoder and rMATS.

(**E, G**) Proportion of genes with functional change and effect scores of ‘SpliceDecoder-enriched’ and ‘rMATS only’ terms for MYC 8h or 24h *vs.* control. ‘SpliceDecoder-enriched’ includes terms with greater significance in SpliceDecoder enrichment compared to rMATS and ‘rMATS only’ includes terms uniquely significant in rMATS enrichment test. Statistical significance was estimated by using 2sample KS test and one-sided Mann-Whitney U test.

(**H**) Decision tree for a group with higher expression of Sim-TX: For each DSE and Ref-TX pair, we consider the type of ‘Simulated event’ and the direction of ‘ΔPSI’ to assign the group with higher expression of Sim-TX. This process assigns the same direction of ‘ΔPSI’ for Inclusion simulation and the opposite direction of ‘ΔPSI’ for skipping simulation.
